# Dynamical modelling of proliferative-invasive plasticity and IFNγ signaling in melanoma reveals mechanisms of PD-L1 expression heterogeneity

**DOI:** 10.1101/2023.01.09.523355

**Authors:** Seemadri Subhadarshini, Sarthak Sahoo, Shibjyoti Debnath, Jason A. Somarelli, Mohit Kumar Jolly

## Abstract

Phenotypic heterogeneity of melanoma cells contributes to drug tolerance, increased metastasis, and immune evasion in patients with progressive disease. Diverse mechanisms have been individually reported to shape extensive intra- and inter-tumoral phenotypic heterogeneity, such as IFNγ signaling and proliferative to invasive transition, but how their crosstalk impacts tumor progression remains largely elusive. Here, we integrate dynamical systems modeling with transcriptomic data analysis at bulk and single-cell levels to investigate underlying mechanisms behind phenotypic heterogeneity in melanoma and its impact on adaptation to targeted therapy and immune checkpoint inhibitors. We construct a minimal core regulatory network involving transcription factors implicated in this process and identify the multiple “attractors” in the phenotypic landscape enabled by this network. Our model predictions about synergistic control of PD-L1 by IFNγ signaling and proliferative to invasive transition were validated experimentally in three melanoma cell lines – MALME3, SK-MEL-5 and A375. We demonstrate that the emergent dynamics of our regulatory network comprising MITF, SOX10, SOX9, JUN and ZEB1 can recapitulate experimental observations about the co-existence of diverse phenotypes (proliferative, neural crest-like, invasive) and reversible cell-state transitions among them, including in response to targeted therapy and immune checkpoint inhibitors. These phenotypes have varied levels of PD-L1, driving heterogeneity in immune-suppression. This heterogeneity in PD-L1 can be aggravated by combinatorial dynamics of these regulators with IFNγ signaling. Our model predictions about changes in proliferative to invasive transition and PD-L1 levels as melanoma cells evade targeted therapy and immune checkpoint inhibitors were validated in multiple data sets from *in vitro* and *in vivo* experiments. Our calibrated dynamical model offers a platform to test combinatorial therapies and provide rational avenues for the treatment of metastatic melanoma. This improved understanding of crosstalk among PD-L1 expression, proliferative to invasive transition and IFNγ signaling can be leveraged to improve the clinical management of therapy-resistant and metastatic melanoma.

## Introduction

Cutaneous malignant melanoma arises from fully differentiated, pigment-producing melanocytes. It is a deadly and highly aggressive form of skin cancer infamous for its high metastatic potential. It carries high mutational burden and is commonly driven via activating mutations in BRAF and NRAS that activate constitutive signaling of mitogen-activated protein kinase (MAPK) (1). The advent of targeted BRAF/MEK inhibitors has been a key breakthrough in the clinical management of advanced melanoma. In 2011, vemurafenib was the first FDA-approved drug to target – BRAF ^V600E^, a key mutation that caused constitutive activation of MAPK pathway and hyper-proliferation (2, 3). However, metastasis in melanoma is largely mediated by phenotypic plasticity and cellular heterogeneity rather than genetic mutation-driven phenomena (4, 5). Melanoma samples exhibit marked intra- and inter-tumor heterogeneity, i.e., they are comprised of diverse phenotypic subpopulations, each with their characteristic molecular profiles and functional attributes of proliferation rate, drug susceptibility and metastatic potential. Reversible bidirectional switching between a proliferative cell state and a slow-cycling invasive state can promote metastasis *in vivo* (5). Such proliferative to invasive transition (PIT) and its reverse invasive to proliferative transition (IPT) is often governed by dynamic alterations in the local microenvironment (6), including therapy-induced adaptive cellular changes (7, 8). Recent characterization of melanoma cell lines and tumor samples at a single-cell level have indicated that phenotypic heterogeneity in melanoma extends beyond the binary proliferative-invasive paradigm, and cells can dynamically acquire a spectrum of phenotypes (9–11). Such plasticity and heterogeneity subsequently facilitate resistance to clinical interventions (12) (chemotherapy, immunotherapy, targeted therapy, *etc.*), posing major challenges in designing effective therapies against advanced melanoma (13).

Melanoma is among the most sensitive malignancies to immune modulation. Immune checkpoint proteins expressed on cancer cell surface such as PD-L1 can binding to inhibitory PD-1 receptors expressed on activated T cells (14). PD-L1 overexpression can accelerate CD4+ T effector cell exhaustion (15) or can diminish CD8+ T cell cytotoxicity (16). High levels of PD-L1 in cancer cells can also directly confer resistance to T cell-mediated death without specifically relying on PD-1-dependent inhibition of T cells (14, 17). In melanoma, PD-L1 levels have been reported as an independent prognostic factor (18). Another breakthrough in treating metastatic melanoma has been the use of immunotherapy. Pembrolizumab was the first FDA-approved PD-L1/PD-1 blocking drug to treat advanced melanoma (19). Importantly, in melanoma cell lines, heterogeneity in PD-L1 expression levels were observed to be independent from any driver mutation in MAPK or PI3K pathway (20, 21). Such heterogeneity in PD-L1 expression profiles has been reported within and between patients (22), and can enable resistance to immunotherapy via non-mutational mechanisms (23). Moreover, recent experiments revealed an association between PD-L1 expression levels and cellular dedifferentiation to an invasive phenotype (21), thus indicating how phenotypic plasticity has the potential to thwart immunotherapy as well. The mechanistic underpinnings driving the emergence of such non-genetic heterogeneity remain largely elusive. Therefore, to better design therapeutic strategies, the dynamics of phenotypic plasticity during proliferative to invasive transition and its impact on PD-L1 expression heterogeneity needs to be better understood.

Here, we have identified a minimal gene regulatory network that can recapitulate the phenotypic heterogeneity along the proliferative-invasive spectrum and its impact on PD-L1 heterogeneity. We mathematically modeled the emergent dynamics of this regulatory network, consisting of key proliferative (MITF, SOX10) and invasive players (JUN, SOX9, ZEB1) and integrating the literature-derived regulatory links among them. Our simulations performed for this network over a parameter ensemble reveal the existence of distinct phenotypes – proliferative, invasive, neural crest-like and intermediate – as an outcome of the network topology. We then couple PD-L1 and one of its key regulators, IFNγ signaling to the network to elucidate the varying immune evasion traits along the proliferative-invasive axis. We observed that neural crest and invasive cells have greater propensities to exhibit high PD-L1 levels with low IFNγ signaling compared to intermediate and proliferative cells. The PD-L1 levels of all the above-mentioned phenotypes can be further enhanced with high IFNγ signaling, which is indicative of their enhanced immune evasion potential. Moreover, cells can reversibly switch from a proliferative/low PD-L1 state to neural crest/high PD-L1 state or invasive/high PD-L1 state, showcasing dynamic changes in PD-L1 levels during dedifferentiation. We quantitatively analyze the coupling between expression levels of PD-L1, IFNγ signaling and proliferative-invasive status of melanoma cells to uncover a mechanistic understanding driving the patterns of PD-L1 heterogeneity in patient-derived melanoma cells prior to and following anti-BRAF targeted therapy. Our model predictions are validated by extensive transcriptomic data analysis at both bulk and single-cell levels for publicly available melanoma RNA-seq data as well as via *in vitro* experiments in proliferative and neural crest-like cell lines. We show that drug resistance to targeted therapies and immune checkpoint inhibitors can occur due to pre-existing cell state heterogeneity or due to drug induced heterogeneity. Finally, we postulate how this improved understanding of crosstalk among PD-L1 expression, proliferative to invasive transition and IFNγ signaling can be leveraged to improve clinical management of therapy-resistant and metastatic melanoma.

## RESULTS

### A core gene regulatory network explains the patterns of non-genetic heterogeneity along the proliferative-invasive axis in melanoma cells

Phenotypic plasticity in cancer cells is driven by a myriad of complex molecular interactions that drive coordinated changes in molecular and functional aspects of various biological pathways. Identifying a minimal regulatory network that can represent many such experimentally reported interactions and whose emergent dynamics can recapitulate the hallmark patterns of phenotypic plasticity and heterogeneity is, therefore, a first important step to elucidate systems-level behavior, while effectively approximating the biological system at hand. To examine the plasticity of melanoma cells along the proliferative-invasive axis, we identified a regulatory network that involves transcription factors implicated in a proliferative to invasive transition – MITF, JUN, ZEB1, SOX10 and SOX9. MITF can drive the proliferative phenotype at least partly through LEF1, and transactivate its own promoter (24, 25). It can also suppress JUN, a regulator of the invasive cell state, through binding to its enhancer (26). Furthermore, Jun can suppress the activity of MITF directly and/or indirectly (26), and self-activate transcriptionally through forming Jun/ATF-2 (27) heterodimers. Thus, MITF and JUN are engaged in mutual antagonism. Similarly, MITF and ZEB1 can form a “toggle switch” (28), such that MITF directly represses ZEB1 (29), and ZEB1 can bind to the MITF promoter region and inhibit MITF (30). Similar to MITF and Jun, ZEB1 can auto-activate via multiple feedback loops (31). SOX10 has been reported in maintaining a proliferative state (32), but it has also been shown to promote invasive features in melanoma (33, 34). SOX10 inactivation leads to SOX9 upregulation, and SOX9 can downregulate SOX10 expression by binding to SOX10 promoter, thus indicating mutual antagonism between them (32). SOX9 can also be involved in upstream enhancer mediated positive autoregulation (35). Finally, SOX10 has been shown to activate the expression of MITF by directly binding to its promoter (36, 37). Together, these interactions constitute a gene regulatory network (GRN) incorporating the complex crosstalk among diverse players of proliferative to invasive transition or its reverse invasive to proliferative transition (**Fig 1A**).

**Figure 1:**
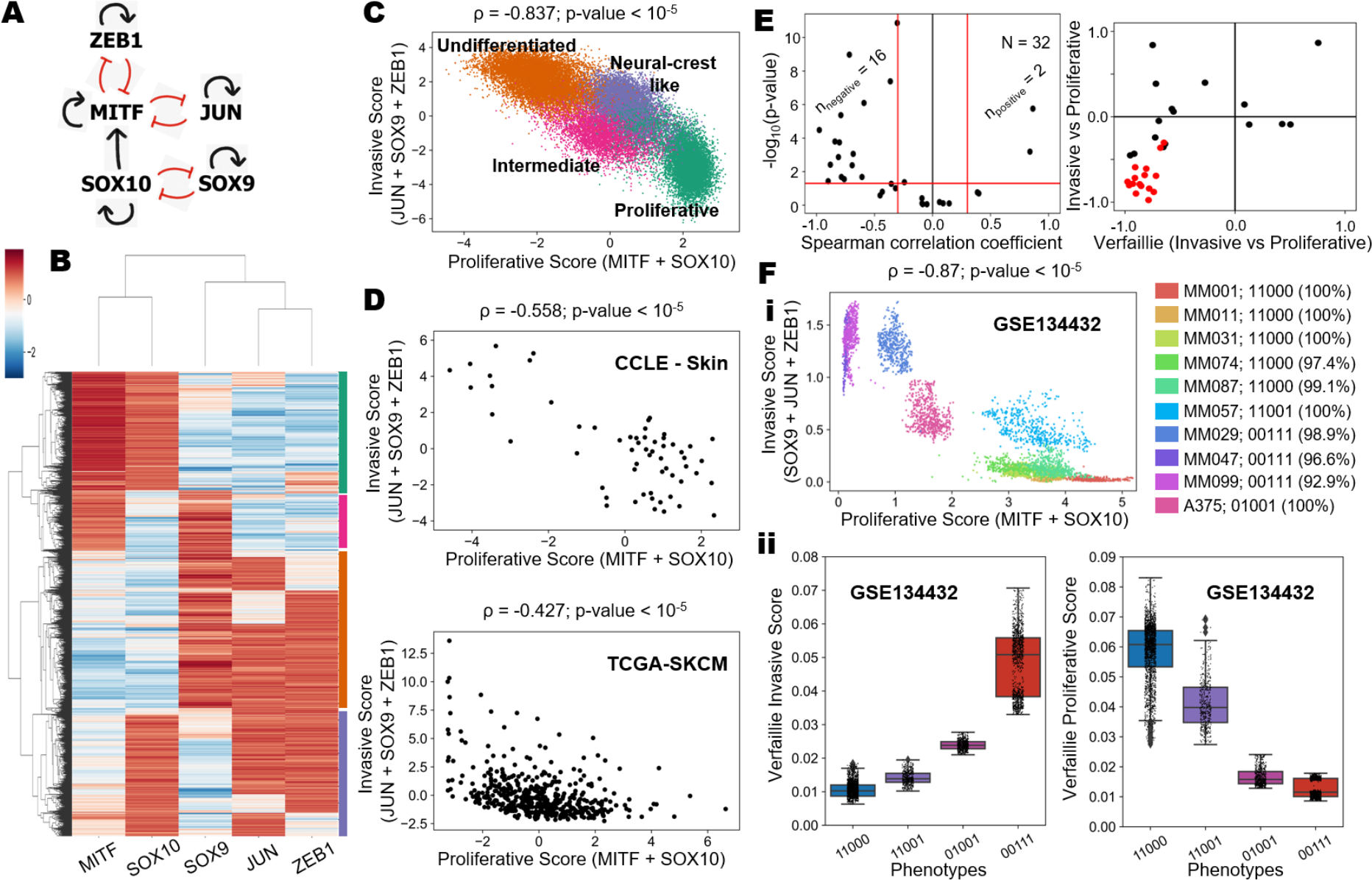
Non-genetic heterogeneity along the proliferative-invasive spectrum enabled by an underlying gene regulatory network in melanoma. A) Gene regulatory network to model the proliferative-invasive heterogeneity in cell states in melanoma. The red hammerheads represent inhibitory links, and the black arrows represent activating connections. B) Hierarchically clustered heatmap of simulated steady states allowed by the gene regulatory network and qualitative classification of the four emergent cell states. C) Scatter plot showing all the steady states projected onto the proliferative - invasive plane defined by sum of expression of MITF, SOX10 (proliferative score) and those of SOX9, ZEB1 and JUN (invasive score). The steady states are colored by cluster labels obtained from hierarchical clustering. D) Scatter plot of the association between the proliferative (MITF+SOX10) and invasive (SOX9+ZEB1+JUN) scores for the CCLE group of skin cancer cell lines (top). Scatter plot showing association between the proliferative (MITF+SOX10) and invasive (SOX9+ZEB1+JUN) scores for clinical samples from TCGA cohort of SKCM patients (bottom). E) Volcano plot for the results of meta-analysis of melanoma datasets, accounting for the associations between the proliferative and invasive scores; nnegative and npositive denote the number of datasets (out of 32) that are correlated negatively (Spearman correlation coefficient < -0.3; p-value < 0.05) and positively (Spearman correlation coefficient > 0.3; p-value < 0.05) (left). Scatter plot for the comparisons of the proliferative-invasive correlations based on the five gene signature and the gold standard Verfaillie signatures. Points colored in red are datasets that show significant correlations in both metrics (right). **F) i)** Scatterplot of single cell RNA-Seq data showing each cell of each cell line projected on the proliferative-invasive plane based on the imputed expression of five modelled master regulators. For each cell line the dominant phenotype is binary and the corresponding abundances have been reported. **ii)** Boxplots of cells categorized by the dominant binary phenotypes based on the five gene signature and their invasive (left) and proliferative (right) ssGSEA scores based on Verfaillie signatures.

We simulated the dynamics of this GRN using the RACIPE framework to identify the steady state expression patterns that represent the “possibility space” of phenotypes enabled by this GRN. The RACIPE framework simulates the dynamics of a GRN through a set of coupled ordinary differential equations (ODEs), with each ODE for a node in the GRN, and kinetic parameters corresponding to production, degradation and regulation being sampled from biologically-relevant ranges (38). The ODEs are then solved across diverse parameter sets and initial conditions to identify the ensemble of steady states. This ensemble is indicative of possible phenotypes allowed by the corresponding network topology. For the GRN considered here, we first plotted the ensemble of steady state solutions as a heatmap (**Fig 1B**). Upon hierarchical clustering, we observed that the invasive genes JUN, ZEB1 and SOX9 were largely co-expressed and so were the proliferative genes MITF and SOX10. Qualitatively speaking, this heatmap revealed four phenotypes – proliferative (MITF^hi^/SOX10^hi^), undifferentiated (SOX9^hi^ with JUN and/or ZEB1 high), neural crest-like (SOX10^hi^ with cJUN and/or ZEB1 high) and intermediate (MITF^hi^/SOX9^hi^) (**Fig S1A**). Co-expression of SOX10 along with invasive markers, such as ZEB1 and/or JUN, has been shown to be necessary to maintain the neural crest-like phenotype with invasive characteristics in development (39, 40). To classify the ensemble of steady states into cellular phenotypes more systematically, we defined two scores: proliferative score (sum of z-normalized gene expression values of proliferative master regulators - MITF and SOX10) and invasive score (sum of z-normalized gene expression values of invasive marker genes - SOX9, JUN and ZEB1). We then projected all the steady state solutions onto a scatter plot of corresponding proliferative and invasive scores (**Fig 1C**). The four phenotypes observed in the heatmap can also be recapitulated as distinct clusters in this proliferative-invasive plane, with the proliferative and invasive scores showing a strong negative correlation (ρ = -0.837; *p* <10^-5^). Similarly, upon coloring the steady states by a “neural crest-like score” (z-normalized expression values of SOX10+JUN+ZEB1), we observed the neural crest-like phenotype present amidst the proliferative-invasive spectrum (**Fig S1B**). These phenotypes identified by our dynamical model are in accordance with previously reported multi-stage differentiation and de-differentiation states, which define melanoma subtypes with the neural crest-like phenotype as an intermediate cell state between the terminal melanocytic proliferative state and the undifferentiated invasive cell state (41).

To cross-validate the observed trend from model simulations, we analyzed a cohort of skin cancer cell lines included in the Cancer Cell Line Encyclopedia (CCLE) (42) (**Fig 1D; top**). In support of the model simulations, we found that proliferative and invasive scores were strongly negatively correlated in the CCLE skin cancer cell line cohort (ρ = -0.558; *p* <10^-5^). This pattern is also reiterated in the primary tumor cohort from The Cancer Genome Atlas (TCGA) (ρ = -0.427; p <10^-5^) (**Fig 1D; bottom**). Furthermore, in both CCLE and TCGA samples, we observed that the proliferative score defined based on MITF and SOX10 levels correlated strongly positively with the enrichment of a much larger gene set defined for proliferative status in melanoma by Verfaillie *et al.* 2015 (43). Similarly, the invasive score derived from the model (defined as the sum of expression of SOX9, JUN and ZEB1) correlated strongly with the enrichment of the Verfaillie gene set used to define invasive status in melanoma (**Fig S1C-D**). These observations strongly suggest that the five genes involved in the GRN model seem sufficient to capture the major patterns in the proliferative and invasive nature of melanoma samples, both for cell lines and patient tumors. As expected, the proliferative and invasive scores defined based on Verfaillie gene expression signatures negatively correlated with each other in both CCLE and TCGA (**Fig S1E**), with the extent of antagonism similar to that seen using our five gene signature (**Fig 1D**).

To ensure the reliability of the chosen five genes for our regulatory network, we compared the correlation of the five identified master regulators (MITF, SOX10 for proliferative; JUN, ZEB1, SOX9 for invasive) against a distribution of correlation coefficients of random combinations of transcription factors (negative control) with the Verfaillie proliferative and invasive signatures in both CCLE skin cancer cell line group and TCGA SKCM patient cohort (**Fig S2**). We observed that the MITF-SOX10 pair and the JUN-ZEB1-SOX9 triplet are generally more strongly correlated to the Verfaillie proliferative and invasive signatures, respectively, compared to any random combination of two or three transcription factors respectively in the CCLE and TCGA datasets, (**Fig S2 A-B**). This suggests that these sets of transcription factors are key drivers of proliferative-invasive phenotypic regulation in melanoma. Furthermore, we found that the trend remains consistent for the MITF-SOX10 pair even when the background reference distribution is created by sampling pairs of transcription factors exclusively from the Verfaillie proliferative gene list (**Fig S2C-D**). However, a similar analysis for the triplet of invasive genes did not show very strong trends and specificity in CCLE and TCGA datasets (**Fig S2C-D**). To improve the specificity of our analysis, we added TCF4 and IRF1 to the signature, as they were among the top correlated transcription factors, which significantly improved the observed trends (**Fig S2E**).

To provide further validation of our observations in the CCLE and TCGA cohorts, we analyzed 32 publicly available melanoma bulk transcriptomic datasets (**Table S1**) and observed a significant negative correlation (ρ < -0.3, *p* <0.05) between the defined proliferative and invasive score as the predominant trend (**Fig 1E; left**). In 27 out of the 32 datasets, our proliferative score defined based on our minimalistic GRN (MITF+SOX10) correlated positively with the Verfaillie proliferative score (**Fig S3A**). Similarly, in 16 out of 32 datasets, our invasive score (=SOX9+ JUN+ ZEB1) correlated positively with Verfaillie invasive score (**Fig S3B**). These results indicate that our minimal GRN is sufficient to capture the proliferative-invasive antagonism, observed across many datasets when compared against the well-established Verfaillie proliferative and invasive signatures (**Fig 1E, Fig S3C**). Finally, we estimated the strength of antagonism between proliferative and invasive scores vis-à-vis that between the Verfaillie proliferative and invasive gene signatures (43). We found that the anti-correlation between the two scores defined based on our GRN is largely consistent (in 16 out of 32 datasets) with the Verfaillie scores defined proliferative-invasive spectrum (**Fig 1E; right**). The antagonism between the proliferative and invasive scores, as well as the positive correlation between the invasive score and Verfaillie invasive signature in both TCGA and CCLE datasets, were also seen when using the updated invasive score (including IRF1, TCF4, SOX9, JUN, ZEB1) (**Fig S3D-E**). The updated invasive score also showed similar trends in meta-analysis of melanoma datasets as noted earlier (compare **Fig S3F, i** with **Fig 1E**). Additionally, 25 out of 32 datasets showed a positive correlation (ρ > 0.3, p <0.05) between the updated five gene invasive score and the Verfaillie invasive score (**Fig S3F, ii**). Thus, our model observations can recapitulate the patterns of phenotypic heterogeneity across many publicly available melanoma datasets, including *in vitro*, *in vivo* and patient sample data sets.

To analyze phenotypic heterogeneity at a single-cell level in melanoma with respect to the five genes in the core GRN, we analyzed single-cell transcriptomic profiles of patient-derived melanoma cultures (GSE134432) along the proliferative-invasive axis (9). We projected 39,263 cells belonging to ten genetically homogenous patient-derived malignant melanoma (MM) lines on the proliferative-invasive plane (**Fig 1F, i**). A density histogram for the expression of the five core genes, fitted with a kernel density estimate, revealed a largely bimodal distribution (**Fig S4A**) just as in our simulations (**Fig S1A**). Upon binarization of gene expression levels in each melanoma cell (see **Methods**), we quantified the gene state as high (1) or low (0). We noticed that a large majority (> 92%) of the cells for specific cell lines can be categorized into either of the four observed phenotypes i.e., the phenotypes of the cells were found to be consistent within their cell line. The cells belonging to the cell lines - MM001, MM011, MM031, MM074 and MM087 - can be assigned to a proliferative subtype with high levels of MITF and SOX10 and low levels of SOX9, JUN and ZEB1 ({MITF, SOX10, SOX9, JUN, ZEB1} = 11000). In contrast, MM047, MM029 and MM099 cell lines are typically invasive, with low levels of MITF and SOX10 and high levels of SOX9, JUN and ZEB1 high ({MITF, SOX10, SOX9, JUN, ZEB1} = 00111). Interestingly, A375 can be classified as a neural crest like cell type with high levels of SOX10 and ZEB1 and low levels of MITF, SOX9 and JUN ({MITF, SOX10, SOX9, JUN, ZEB1} = 01001). Moreover, the cell line MM057 belonged to an intermediate phenotype with a subset of both proliferative and invasive factors being relatively high. This cell state can also be found as a subset of the proliferative phenotype in our simulation results (**Fig 1B**), further underscoring the applicability of a parameter ensemble modelling approach to capture the “possible cell states” allowed by a gene regulatory network. Similar patterns were revealed when cell lines were projected in a 2-D proliferative and invasive score plane (**Fig S4B**). After the classification of these cell-lines into various phenotypes along the proliferative-invasive spectrum based on the five gene regulatory network model, we tested our classification against the activity-based score of Verfaillie proliferative and invasive signatures (**Fig 1F, ii**). The phenotype corresponding to a high expression of MITF and SOX10 and low expression of SOX9, JUN and ZEB1 (11000) had the highest Verfaillie proliferative score and the least Verfaillie invasive score. Conversely, the 00111 phenotype where the invasive players are more highly expressed compared to the proliferative ones had low Verfaillie proliferative scores and high Verfaillie invasive scores. The other phenotypes where one or more of both proliferative and invasive genes are high were ranked as intermediate with respect to the extreme 00111 and 11000 phenotypes, marking the ends of the Verfaillie proliferative-invasive axis. Similarly, the cells we categorized as a neural crest-like phenotype based on MITF, SOX10, SOX9, JUN, ZEB1 ({MITF, SOX10, SOX9, JUN, ZEB1} = 01001) had enrichment of activity of the neural crest associated gene signature (**Fig S4C**).

Overall, we demonstrate that emergent dynamics of a minimal five-gene-based GRN can explain the hallmark patterns underpinning phenotypic heterogeneity in melanoma cell lines and tumors, both at the bulk and single-cell transcriptomic levels, along the proliferative-invasive spectrum.

### Perturbations to the gene regulatory network can cause non-genetic phenotypic changes along the proliferative-invasive spectrum during melanoma disease progression

To understand how perturbing our core GRN can impact proliferative-invasive characteristics, we simulated two scenarios: a) downregulation (DE) of SOX10 (**Fig 2A**) and b) over-expression (OE) of SOX9 (**Fig 2E).** Downregulation of SOX10 led to an increase in the frequency of the invasive phenotype and a decrease in the proliferative phenotype, when compared to the unperturbed wild type (WT) network. We also found a decreased frequency of the neural-crest phenotype and an increased proportion of the intermediate phenotype (**Fig 2A**). The model simulations are substantiated by *in vitro* bulk transcriptomic analysis of the effects of SOX10 shRNA-mediated knockdown in M010817 human melanoma cells (GSE37059) (32). SOX10 knockdown led to a decrease in levels of the MITF and SOX10-based proliferative score, indicative of an enrichment of the invasive state (**Fig 2B, left**). Simultaneously, these cells have a higher invasive score, as defined by expression levels of SOX9, JUN and ZEB1 (**Fig 2B**, **right)** as well as with an invasive score defined by the expression of SOX9, JUN, ZEB1, IRF1 and TCF4 (**Fig S5A**). Furthermore, cells with SOX10 knockdown displayed perturbation of the regulon-based proliferative and invasive scores as compared to control cells (**Fig S5A**). We next interrogated the impacts of SOX10 knockdown in the A375 neural crest-like melanoma cell line (GSE180568). Similar to M010817 melanoma cells (**Fig 2B**), SOX10 knockdown in A375 cells led to an increase in the invasive score and a slight decrease in the proliferative scores (**Fig 2C, S5B**). We also found that the increased invasiveness was accompanied by a loss of the neural crest-like signature, indicative of a transition from a neural crest-like phenotype to an undifferentiated phenotype, as predicted by the *in silico* perturbation simulations (**Fig 2A)**. To assess the dynamic trajectory of transition from a more proliferative to a more invasive phenotype, we next analyzed single cell RNA-seq data of SOX10 knock down time-course experiments of three available cell lines (MM057, MM074 and MM087), all of which exhibit a proliferative phenotype (**Fig 1F, i**). Upon projecting the MM057 control and SOX10-siRNA treated cells (GSE134432) on the proliferative/invasive plane, we observed a gradual shift of cells from proliferative to invasive. By 72 hours of SOX10 knockdown MM057 cells were as invasive as the canonical invasive cell lines (MM047, MM099 or MM029), with low levels of MITF and SOX10 and high levels of SOX9, JUN and ZEB1 high (00111) (compare **Fig 2D** with **Fig 1F, i**). Similar trajectories and shifts in phenotypic distributions to varying degrees were observed for MM074 and MM087 cell lines upon knock down of SOX10 (**Fig S5C**).

**Figure 2:**
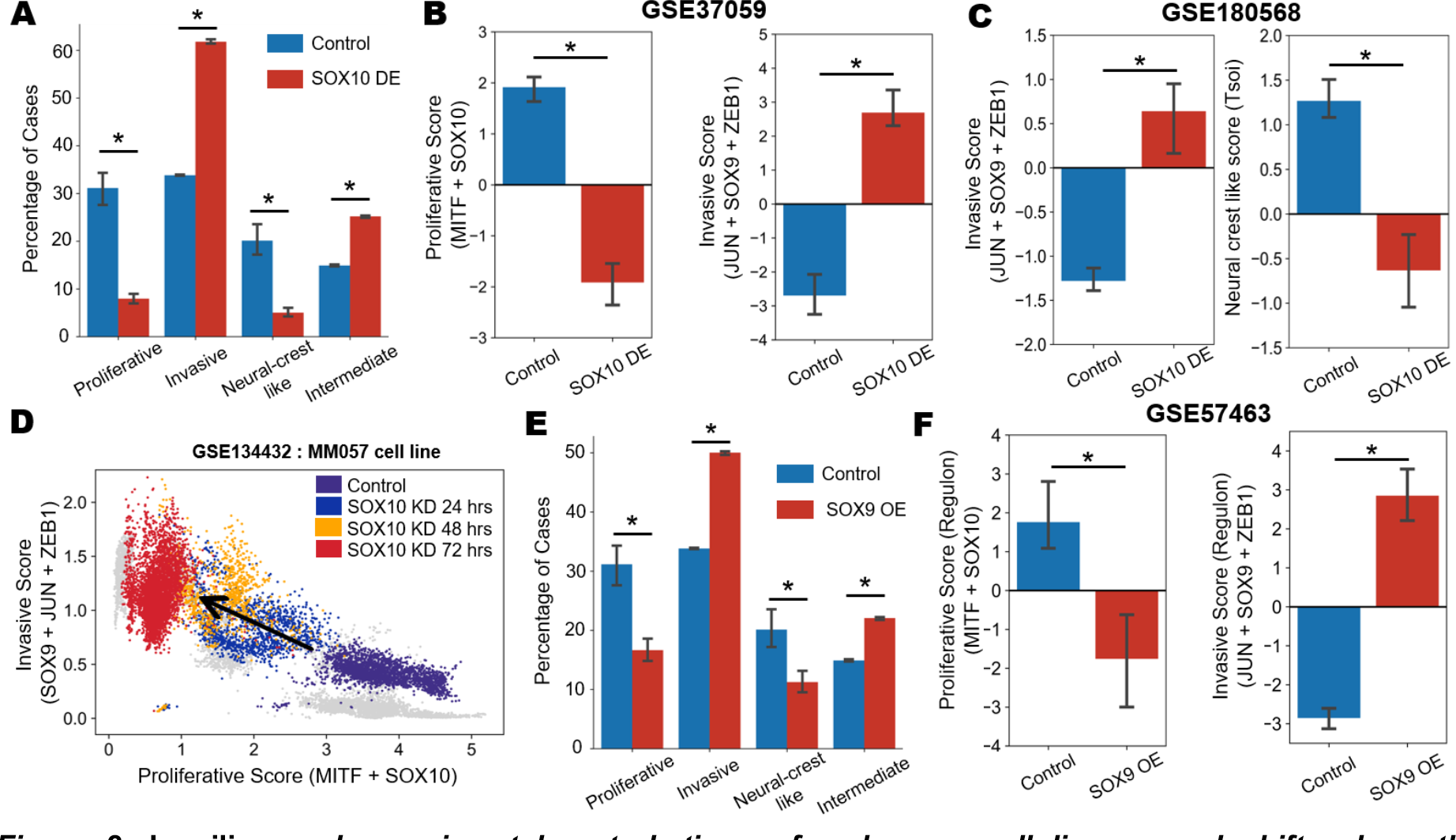
In silico and experimental perturbations of melanoma cell lines reveal shifts along the proliferative-invasive axis. A) Bar plots showing proportions of steady state solutions in control cases compared to SOX10 downregulation across the four phenotypes. Error bars represent standard deviations of n=3 technical RACIPE replicates. **B)** Bar plots of the experimentally observed significant changes (demarcated by *) in the proliferative (defined as the sum of z-normalized expression of MITF and SOX10) (left) and invasive scores (defined as the sum of z-normalized expression of JUN, ZEB1 and SOX9) (right) upon SOX10 knockdown (GSE37059). **C)** Bar plots showing experimentally observed significant changes in the Invasive score (defined as the sum of z-normalized expression of JUN, ZEB1 and SOX9) (left) and neural crest-like ssGSEA score (Tsoi et al.) (right) (GSE180568). **D)** Scatter plot of single cell RNA-Seq data for the transition of SOX10 knockdown cells in MM057 cells at 24h, 48h and 72h in comparison to control data (GSE134432). **E)** Bar plots showing proportions of steady state solutions in control cases compared to SXO9 over expression across the four phenotypes. Error bars represent standard deviations of n=3 technical RACIPE replicates. **F)** Bar plots of the experimentally-observed changes in the proliferative (defined as the sum of z-normalized ssGSEA scores of MITF and SOX10 regulons) (left) and invasive scores (defined as the sum of z-normalized ssGSEA scores of JUN, ZEB1 and SOX9 regulons) upon SOX9 over expression (right) (GSE57463). * represents a statistically significant difference in the levels based on Student’s t-test.

Next, we simulated SOX9 over-expression, which led to an increased frequency of the invasive phenotype (**Fig 2E**). Concurrently, both the proliferative and neural crest-like phenotypes experienced a decrease in their percentage distribution, with a mild increase in that of the intermediate phenotype. To validate this model prediction, we analyzed publicly-available bulk transcriptomic profiles of M010817 human melanoma cells in which SOX9 has been overexpressed (GSE57463) (44). Although we did not observe a marked change in the proliferative score based on the expression of MITF and SOX10 alone (**Fig S5D**), we found a decrease in the proliferative nature (**Fig 2D**, **left**), defined by the activity score of the MITF and SOX10 regulon and an increase in invasiveness (**Fig 2D**, **right**), as defined by the activity score of SOX9, JUN, and ZEB1. The invasive score (JUN, SOX9 and ZEB1 expression) showed a significant increase as reflected in the activity scores of the regulons of the corresponding genes (**Fig S5D**), supporting our model prediction. A similar trend of increasing in invasive score upon SOX9 over expression was also recapitulated using the updated invasive score (**Fig S5D**). Together, these analyses reveal that perturbations in levels of various nodes in our minimal GRN is capable of recapitulating phenotypic switching along the proliferative-invasive axis in melanoma.

### Phenotypic plasticity along the proliferative-invasive axis can alter PD-L1 levels in melanoma cells

After validating the ability of our GRN to predict phenotypic plasticity in melanoma cells, we sought to investigate inter-connection between phenotypic plasticity along the proliferative-invasive spectrum and the immunosuppressive traits of melanoma cells. To understand immune evasion in melanoma, we coupled immune-effector molecules: PD-L1 (suppressor on adaptive immune system (16)) and IFNγ signaling (**Fig 3A**). We extended our GRN based on following experimental observations: 1) IFNγ is a key regulator of PD-L1 levels (21), 2) IFNγ regulates differentiation state of melanoma cells (21, 45, 46), 3) IFNγ can transcriptionally activate PD-L1 via NF-κB and/or JAK-STAT-IRF1 pathway (47, 48), 4) IFNγ can suppress MITF expression by inducing CBR and STAT1 association, thereby inhibiting CREB binding to MITF promoter (49). Likewise, PD-L1 is connected to the GRN through the following experimentally observed connections: 1) MITF can degrade PD- L1 via SA-49 (50), 2) JUN activates the expression of PD-L1 through STAT3 (51), 3) and ZEB1 can upregulate PD-L1 expression (52) and promote immune escape in melanoma (53). Furthermore, we created a variant version of the regulatory network including IRF1 (**Fig S6A**). IRF1 is known to be a potent activator of PD-L1 (54) and is also known to be upregulated by IFNγ signaling (55). SOX10 can also down regulate IRF1 in the context of melanoma cells (56). Thus, PD-L1 and IFNγ signaling crosstalk extensively with multiple drivers of proliferative and invasive phenotypes in melanoma.

**Figure 3:**
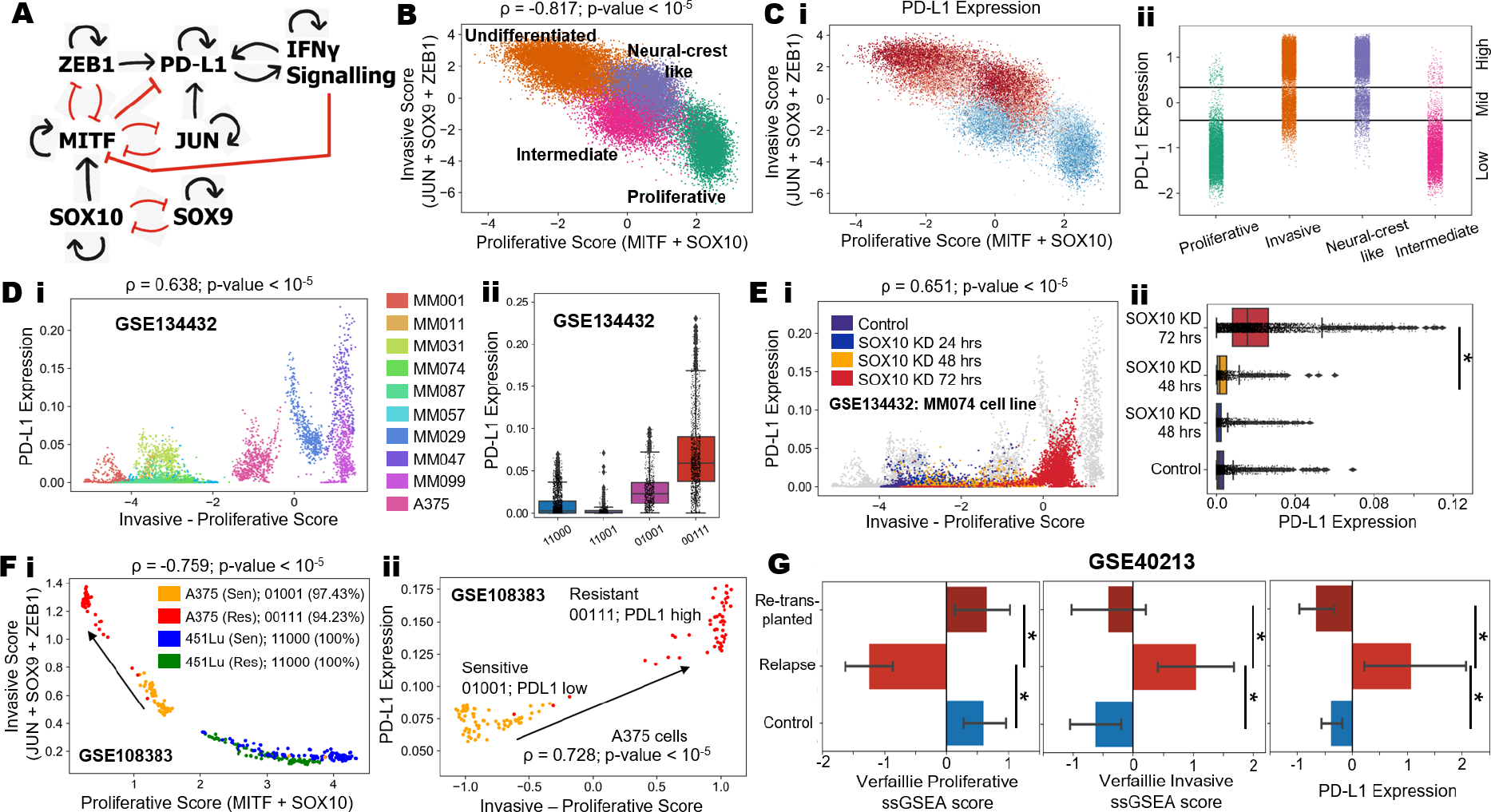
Steady state dynamics of emergent cellular phenotypes upon addition of the PD-L1-IFNγ signaling motif to the gene regulatory network. A) Gene regulatory network coupling the proliferative-invasive regulators with the PD-L1-IFNγ signaling motif in melanoma. The red hammerheads represent inhibitory links, and the black arrows represent activations. B) Scatter plot showing all the steady states projected onto the proliferative - invasive plane defined by expression of MITF, SOX10 (proliferative score) and SOX9, ZEB1 and JUN (invasive score). The steady states are colored by cluster labels obtained from hierarchical clustering; C) i) Scatter plot showing all the steady states projected onto the proliferative - invasive plane colored based on PD-L1 levels. ii) Strip plot showing the PD-L1 steady state levels for the four phenotypes. The horizontal lines mark the stratification of PD-L1 levels into low, mid and high regions. D i) Scatterplot of single cell RNA-Seq data showing each cell of each cell line projected on the invasive-proliferative score and the imputed PD-L1 expression plane. ii) Boxplots of cells categorized by the dominant binary phenotypes based on the five gene signature and their imputed PD-L1 levels. E) i) Scatter plot showing increase in PD-L1 levels of SOX10 knockdown cells in MM074 cell line as they transition from a proliferative phenotype to an invasive phenotype along 24h, 48h and 72h time course single cell RNA-seq data in comparison to control data (GSE134432). ii) Boxplots of cells categorized by different timepoints quantifying their imputed PD-L1 levels. F) i) Scatterplot of single cell RNA-seq data projecting cells of two cell lines – A375 (red and orange corresponding to the resistant and sensitive clones, respectively) and 451Lu (green and blue corresponding to the resistant and sensitive clones, respectively) on the proliferative-invasive plane and ii) Scatterplot of single cell RNA-seq data projecting Vemurafenib sensitive and resistant A375 clones with the difference of invasive and proliferative scores on the x-axis and the PD-L1 levels on the y-axis. G) Barplot showing the proliferative scores (left), invasive scores (middle) and PD-L1 levels (right) across control cells, relapsed cells, and cells retransplanted into mice after relapse (GSE40213). * represents a statistically significant difference in the levels based on Student’s t-test.

As before, we used RACIPE to generate the ensemble of steady-state solutions, enabled by this extended GRN. We plotted all the resultant steady state values on the proliferative-invasive plane after performing hierarchical clustering (**Fig 3B**). We observed similar phenotypic clusters as observed for the earlier GRN (**Fig 1C**), demonstrating a strong proliferative-invasive antagonism (ρ = -0.817; *p* <10^-5^). Upon overlaying PD-L1 levels on the proliferative-invasive plane, we noted that undifferentiated/invasive and neural crest subtypes tended to have high PD-L1 expression whereas intermediate and proliferative cell types had relatively low levels of PD-L1 (**Fig 3C, i**). To quantify the association of PD-L1 for these phenotypes, we first evaluated the histogram of PD-L1 distribution, which showed distinct trimodally (**Fig S6B**). Further analysis revealed that neural crest and undifferentiated states express high to intermediate levels of PD-L1, in contrast to proliferative and intermediate phenotypes, for which PD-L1 expression is primarily in the low range (**Fig 3C, ii**). The variant network including IRF1 (**Fig S6A**) also showed very similar phenotypic clusters (**Fig S6C**) as observed for the network shown in **Fig 3A**.

To validate these model predictions for association of proliferative-invasive status with PD-L1 levels, we investigated the CCLE group of skin cancer cell lines. We used two different metrics to quantify the proliferative-invasive status: a) the five genes included in the GRN (= SOX9 + JUN + ZEB1 – SOX10 – MITF), and b) the Verfaillie gene lists (= ssGSEA (Verfaillie invasive geneset) – ssGSEA (Verfaillie proliferative geneset). In CCLE, we observed weak positive associations of PD- L1 expression with both of these metrics (**Fig S6D**). Similar observations were seen for TCGA cohort of SKCM patients (**Fig S6E**). These trends remained consistent even with the new metric that included updated invasive score including IRF1 and TCF4 (**Fig S6F**). Further, single cell RNA- seq analysis of melanoma cell lines revealed that more invasive cell lines were more likely to express higher levels of PD-L1 compared to the more proliferative cell lines (**Fig 3D, Fig S7A**).

Further, among invasive cell lines, we observed PD-L1 expression levels to be quite heterogenous with cells exhibiting low, medium, and high levels (**Fig 3D, i**). This pattern is reminiscence of our model simulations where the neural-crest like and invasive phenotypes expressed low, intermediate as well as high levels of PD-L1, while the proliferative phenotype had predominantly low PD-L1 levels (**Fig 3C, ii**).

Next, we examined if induction of an invasive state via perturbations to the GRN caused concomitant changes in PD-L1 expression levels. First, we analyzed single cell RNA-seq datasets for siRNA-mediated knockdown of SOX10 in MM057, MM074 and MM087 melanoma lines (**Fig 3E**). PD-L1 levels significantly correlated to the proliferative-invasive status of MM074 cells (**Fig 3E, i**), where PD-L1 expression levels were markedly elevated 72 hours after SOX10 knockdown as compared to the non-transfected controls (**Fig 3E, ii**). Similar trends, albeit weaker, were noted in MM057 and MM087 cells (**Fig S7B**), suggesting a “semi-independent” behavior between PD-L1 expression and the proliferative-invasive status. This pattern denotes that while proliferative to invasive transition alters PD-L1 expression, these levels can be modified by other signaling pathways.

Proliferative to invasive transition in melanoma cells is also often associated with resistance to targeted therapies. More invasive cells are more resistant to targeted therapies such as BRAF/MEK inhibitors (13, 57). Based on this association, we hypothesized that targeted therapy may lead to changes in the GRN that would impact levels of PD-L1. To provide support for this hypothesis, we analyzed single cell RNA-seq data from paired parental and vemurafenib-resistant clones of A375 and 451Lu melanoma cells (GSE108383) (**Fig 3F, i-ii**). Upon projecting the two cell lines onto a proliferative-invasive plane based on the five genes in the GRN, we observed that the A375 cell line was comparatively more invasive in nature compared to the 451Lu cell line. The parental 451Lu cell line was predominantly proliferative based on the five gene signature ({MITF, SOX10, SOX9, JUN, ZEB1} = 11000) while the A375 cell line was predominantly neural crest-like ({MITF, SOX10, SOX9, JUN, ZEB1} = 01001). The vemurafenib-resistant A375 clone was distinctively more invasive compared to its parental counterpart, with decreased SOX10 and increased ZEB1 levels, suggesting that emergence of adaptive resistance to vemurafenib may involve a transition from a neural crest to invasive phenotype in melanoma (**Fig 3F i, Fig S7C**). Conversely, vemurafenib-resistant 451Lu cells did not show such a distinctive phenotypic switch, as the parental and the resistant clones were both proliferative based on the five gene signature. The phenotypic switch from neural crest-like to invasive in A375 cells was accompanied by an increase in PD-L1 levels (**Fig 3F, ii**). No such change was observed in 451Lu cells that did not undergo a switch in phenotypes during the evolution of resistance to vemurafenib (**Fig S7D**). These trends reinforce our model predictions that a switch along the proliferative-invasive spectrum can contribute to increased PD-L1 expression.

We next analyzed transcriptomic profiles for control, relapsed and re-transplanted relapsed tumors for the HCmel3 mouse melanoma model (GSE40213) (**Fig 3G**). We found that the control and the retransplanted tumor samples were more proliferative and less invasive compared to the relapsed melanoma tumors. The levels of PD-L1 levels mirrored the invasive nature of the tumor samples. More importantly, it shows that the increase in PD-L1 levels was reversible as melanoma cells underwent invasive to proliferative transition (**Fig 3G**). Inflammation-induced reversible dedifferentiation has been reported to cause melanoma cells to resist T-cell therapy (8). Overall, our results show that PD-L1 expression in melanoma cells is reversibly associated, at least in part, with the gain of a more invasive phenotype, thereby providing further support for the role of phenotypic plasticity along the proliferative-invasive axis in mediating upregulation of PD-L1.

### Role of IFNγ signaling in induction of PD-L1 levels and dedifferentiation of melanoma cells

A positive association of PD-L1 and IFNγ has been extensively reported experimentally (58–60). We tested whether our model could capture this association. Our simulation results suggest that phenotypes with high or intermediate IFNγ signaling are likely to have high PD-L1 expression, thus highlighting a strong positive influence of IFNγ on PD-L1 (ρ = 0.75; *p* <10^-5^) (**Fig 4A**). To get further supporting evidence for this trend, we analyzed bulk transcriptomic data of melanoma cell lines treated with IFNγ. We found that upon treatment, both the Hallmark IFNγ response pathway activity and PD-L1 expression was enriched, indicating an association between IFNγ signaling and PD-L1 levels (**Fig 4B**). As expected, the PD-L1 expression levels were correlated positively with IFNγ activity levels both in CCLE and TCGA samples (**Fig 4C, Fig S8A**).

**Figure 4:**
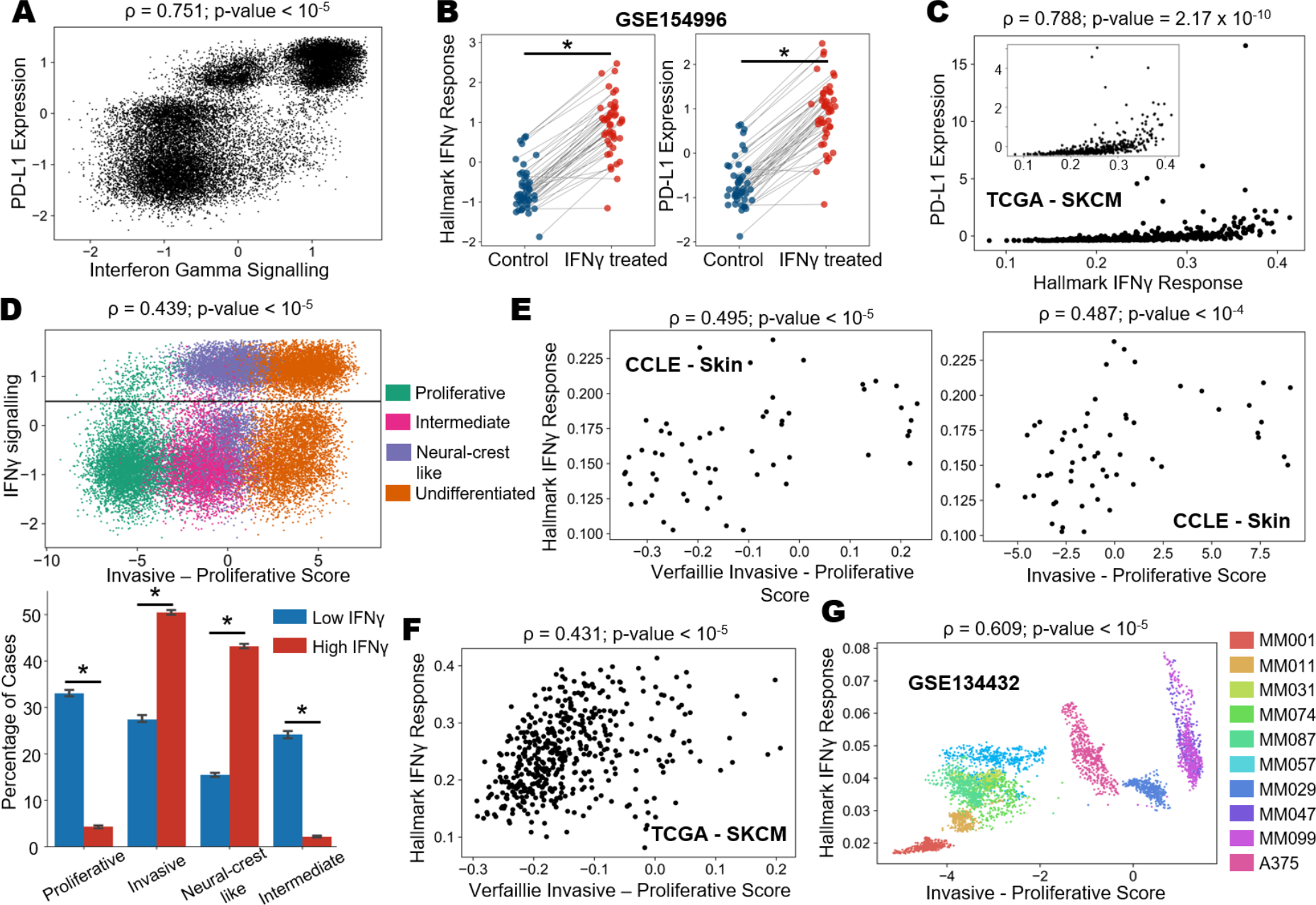
Control of PD-L1 levels and the differentiation state of melanoma cells by IFNγ signaling. A) Scatterplot showing the association between PD-L1 levels and IFNγ signaling at steady state levels. **B)** Paired plot showing changes in levels of **i)** Hallmark IFNγ signaling and **ii)** PD-L1 levels upon treatment of melanoma cells with IFNγ. * represents a statistically significant difference in the levels based on a paired Student’s t-test. **C)** Scatter plots showing association between Hallmark IFNγ response (x-axis) and PD-L1 expression (y-axis) in the TCGA melanoma cohort. **D) (top)** Scatterplot showing the spread of steady state solutions with invasive – proliferative score on the x-axis and IFNγ signaling on the y-axis. The steady states have been colored by the corresponding four discretized phenotypes. The horizontal line represents the stratification between low and high IFNγ signaling regions. **(bottom)** Bar plots showing the fraction of steady state solutions that belong to each of the four phenotypes segregated based on the level of IFNγ signaling. The error bars represent the standard deviation based on n=3 RACIPE replicates. *represents a statistically-significant difference in the levels based on a Student’s t-test. **E)** Scatterplot showing associations between the i) Verfaillie Invasive – Proliferative scores and ii) five-gene signature based invasive – proliferative scores with Hallmark IFNγ signaling. **F)** Scatterplot showing associations between the Verfaillie Invasive – Proliferative scores with Hallmark IFNγ signaling in the TCGA melanoma cohort. **G)** Scatterplot of single cell RNA-Seq data showing individual cells from each cell line projected on the invasive-proliferative and Hallmark IFNγ signaling planes. Each color represents a different cell line.

Next, we investigated how proliferative to invasive transition status associates with IFNγ signaling. Our simulations suggest that, similar to PD-L1, IFNγ signaling is higher in neural crest-like and undifferentiated phenotypes (**Fig S8B, S8C**). Furthermore, while all four phenotypes are observed at lower IFNγ levels, neural crest-like and undifferentiated phenotypes predominate at higher levels of IFNγ (**Fig 4D**). This trend is reminiscent of experimental data suggesting that IFNγ signaling can induce dedifferentiation in melanoma (46). In CCLE and TCGA samples, we observed strong positive associations between the proliferative-invasive status and IFNγ signaling for both the reduced five-gene signature or Verfaillie-based metric of proliferative-invasive status (**Fig 4E, 4F**). This positive association was also recapitulated when considering the updated invasive score and IFNγ signaling (**Fig S8D**). Additionally, treatment of melanoma cells with IFNγ signaling led to significant, but variable decreases in proliferative scores and an increase in invasive scores (**Fig S8E**). Similar results were observed when we analyzed a group of n=8 melanoma cell lines treated with either TNF or IFNγ (**Fig S8F-H**) (GSE152755) (46). While treatment of TNF and IFNγ caused expected significant increases in the corresponding pathways, PD-L1 levels were also induced, especially for IFNγ-treated cells (**Fig S8F**). However, the extent of dedifferentiation in either case was variable (**Fig S8G-H**). This is consistence with reports where transcription factors such as Jun have been shown to have a relatively modest role in IFNγ induced expression of PD-L1 in melanoma cells (48). Analysis of single cell RNA-seq data of ten patient-derived malignant melanoma (MM) lines (GSE134432), revealed a strong positive correlation between intrinsic IFNγ signaling levels and invasive/proliferative scores (p = 0.609; p < 10^-5^) (**Fig 4G**). Together, these results show that increased IFNγ signaling in melanoma can affect levels of PD-L1 and/or differentiation status of melanoma cells via the underlying gene regulatory network.

### Combinatorial control of PD-L1 levels by IFNγ signaling and proliferative-invasive status of cells underlies heterogeneous expression patterns of PD-L1 in melanoma

Our model simulations indicated that proliferative and intermediate states typically have lower PD- L1 levels, but neural crest-like and undifferentiated phenotypes have either intermediate or high levels of PD-L1 (**Fig 5A, left**). The highest levels of PD-L1 observed were also associated with a higher activation of IFNγ signaling (**Fig 5A, right**). These observations suggested that PD-L1 levels could be upregulated through two possible paths: a) increase in invasive nature, and b) enhanced IFNγ signaling. We further quantified the contribution of these two paths and their interdependency.

**Figure 5:**
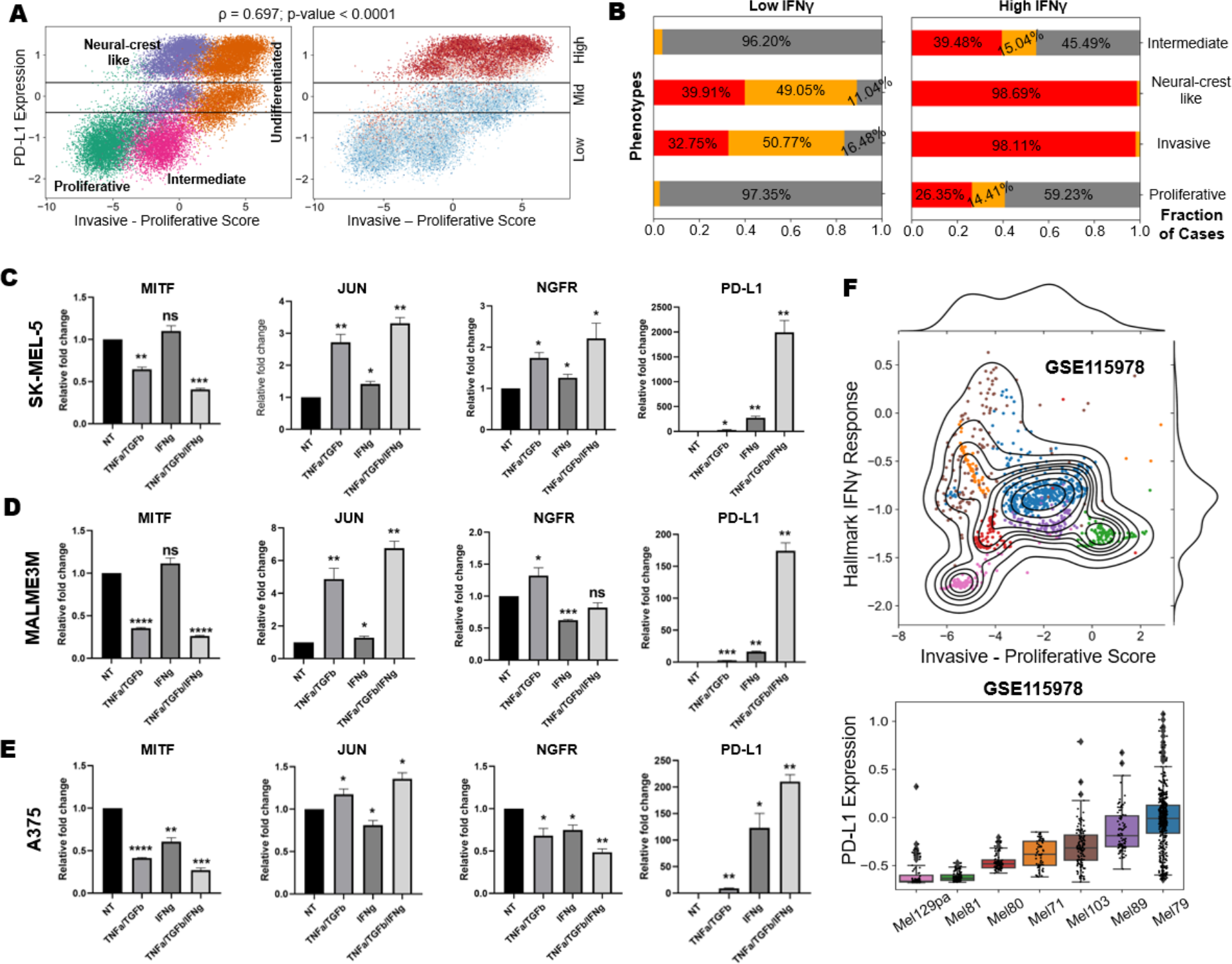
Analysis of coupled dynamics for PD-L1, proliferative-invasive phenotype, and IFNγ signaling. **A)** Associations between invasive – proliferative score and PD-L1 levels colored by the four phenotypes (**left**) and IFNγ signaling levels (**right**). **B)** Stacked bar plots showing the fraction of steady states belonging to low (gray), mid (orange), or high (red) PD-L1 levels in either low IFNγ signaling levels (**left**) or high IFNγ signaling levels (**right**). The main proportion of steady states belonging to each group are indicated. **C, D, E**) Bar plots showing the relative fold changes in the mRNA expression levels of MITF, JUN, NGFR and PD-L1 after 48 hours in non-treated (NT), TNFα/TGFβ treated, IFNγ treated and TNFα/TGFβ /IFNγ treatment conditions in melanoma cells **C)** SK-MEL- 5 **D)** MALME-3M **E)** A375. * (p-value < 0.05), ** (p-value < 0.01), *** (p-value < 0.001), **** (p-value <0.0001) represents a statistically significant difference in the levels based on paired t-test. **F)** Scatterplot of single cells from melanoma patients before immune checkpoint inhibitors therapy projected on the Invasive-Proliferative score and Hallmark IFNγ signaling (**top**) and box plots of their corresponding PD-L1 levels in a patient specific grouping sorted ascending order according to median expression values (**bottom**). Each color in scatterplot corresponds to a particular patient sample.

Thus, we segregated the scenarios belonging to high vs. low IFNγ signaling and subsequently calculated the conditional probability of each of the four phenotypes to display low, medium, and high levels of PD-L1. For low IFNγ signaling, proliferative and intermediate phenotypes were most likely (> 95%) to express low PD-L1 levels, but for neural crest-like and invasive phenotypes, PD- L1 levels were either medium (49% - 51%) or high (32% - 40%) (**Fig 5B**, **left**). Importantly, while proliferative and intermediate phenotypes were more homogeneous in terms of PD-L1 levels, the neural crest-like and invasive ones exhibited higher variance. This behavior recapitulates the trends seen in a single-cell RNA-seq cohort of melanoma cell lines spread across the proliferative-invasive spectrum (**Fig 3D, i**). Together, these results suggest that in absence of IFNγ signaling, a proliferative to invasive transition can increase PD-L1 levels. For high IFNγ signaling, amplified heterogeneity in PD-L1 is seen for proliferative and intermediate phenotypes. Conversely, invasive and neural crest-like are most likely (> 98%) to display high PD-L1 and thus more homogenous (**Fig 5B**, **right**). Importantly, we notice that a substantial proportion of cells (>25%), irrespective of their proliferative-invasive status, can exhibit high PD-L1 levels, under scenarios of high IFNγ signaling (compare the proportion of PD-L1 high cells (red) across phenotypes in **Fig 5B**, right vs. left). This trend is consistent with experimental reports showing proliferative to invasive transition inducing factors such as JUN having a relatively minor influence on IFNγ-induced PD-L1 expression (48). Put together, our simulations highlight that increased IFNγ signaling, and a more invasive nature can both augment PD-L1 levels.

To experimentally test these model predictions, we chose three melanoma cell lines – SK-MEL-5, MALME3M (both representative of a proliferative phenotype) and A375 (representative of the neural-crest phenotype) based on their gene expression profiles in the CCLE group of melanoma cell lines. SK-MEL-5 and MALME3M have higher proliferative scores compared to their invasive scores and lie towards the proliferative end of the spectrum on a two-dimensional proliferative-invasive plane of the Verfaillie gene sets (**Fig S9A**). A375 cells, however, are more towards the center of this phenotypic spectrum (**Fig S9A, left**) and exhibit higher levels of neural crest markers compared to MALME3M and SK-MEL-5 (**Fig S9A, right**). To test our model predictions, we treated each of these cell lines with either a combination of TNFα and TGFβ to induce a proliferative to invasive transition (PIT) or we treated the cell lines with IFNγ as an independent modulator of PD- L1 levels and assessed the mRNA levels of MITF (proliferative marker), JUN (invasive marker), NGFR (neural-crest marker) and PD-L1 levels after a duration of 48 hours. Treatment of MALME3M and SK-MEL-5 cells with both TNFα and TGFβ led to significantly lower levels of MITF and higher levels of JUN, indicative of a less proliferative and more invasive cell state (**Fig 5C-D**). These changes were also accompanied by significant increase in NGFR levels, indicating the transition from a proliferative to a neural crest/invasive phenotype, as well as enhanced PD-L1 levels (**Fig 5C-D, right**). Treatment with IFNγ alone, on the other hand, did not cause such drastic changes for proliferative (MITF), invasive (JUN) and neural crest (NGFR) markers (**Fig 5C-D**). One explanation for this modest impact of IFNγ alone on these markers can be its epigenetic mode of action, which are often discernible over longer time scales (61). Despite minimal impact on proliferative to invasive transition, IFNγ treatment alone induced PD-L1 levels to an even higher extent as compared to TNFα/TGFβ treatment (**Fig 5C-D, right**), thus demonstrating that IFNγ can induce PD-L1 levels efficiently with minimal effects on proliferative to invasive transition at relatively short timescales. Upon a combinatorial treatment of TNFα/TGFβ and IFNγ, PD-L1 levels increased synergistically in both proliferative cell lines, potentially showing an emergent relationship between the two modes of induction of PD-L1 levels (**Fig 5C-D, right**). This observation resonates well with our simulation results that under conditions of low IFNγ levels, proliferative cells are predominantly (97.35%; **Fig 5B, left**) low in terms of PD-L1 levels, but under scenarios of high IFNγ levels and a switch (induced by TNFα/TGFβ) to more neural crest like or undifferentiated phenotype, most cells (> 98.1%; **Fig 5B, right**) express high PD-L1.

Next, we checked the effects of individual and combinatorial treatments of TNFα/TGFβ and IFNγ on a neural crest cell line, A375. We observed a robust downregulation of proliferative (MITF) and upregulation of invasive (JUN) markers upon TNFα/TGFβ treatment (**Fig 5E).** IFNγ treatment alone inhibited MITF, but had modest effects on levels of JUN, consistent with earlier reports (62). Earlier observations report that neural-crest phenotype has the highest NGFR expression (63) and thus we would expect that a transition to a more undifferentiated phenotype would lead to a decrease in NGFR levels. Validating this hypothesis, we observed robust downregulation of NGFR mRNA expression (**Fig 5E**), indicating that the cells that were originally of the neural crest phenotype transitioned to a more undifferentiated invasive phenotype. This was again accompanied by robust upregulation of PD-L1 levels in TNFα/TGFβ and IFNγ treated cases with the combinatorial treatment (**Fig 5E, right**), showing the highest increase as predicted by our model.

Finally, to determine if these trends were observed in melanoma cells from treatment naïve patients (GSE115978) (64), we assessed how PD-L1 levels depended on both the extent of proliferative to invasive transition and activity of IFNγ signaling. Typically, cells from each patient sample cluster together in this plot of invasive score and IFNγ signaling activity (**Fig 5F, top**), showcasing greater inter-patient phenotypic variability than intra-patient variability. Across the patient samples, we observe scenarios of varying degrees of invasive score and IFNγ signaling, enabling us to evaluate how these both axes control PD-L1 levels.

First, we focused on pre-treatment samples. Compared to other cells, Mel129pa cells (pink cluster) were less invasive and had lower IFNγ signaling (**Fig 5F, top**). These cells had the lowest PD-L1 levels (**Fig 5F, bottom**), thus validating our model predictions that proliferative phenotype with low IFNγ signaling has minimal PD-L1 expression (**Fig 5B, left**). These analyses support a model in which at least one of the two axes is needed to increase PD-L1 levels: proliferative-invasive transition or IFNγ signaling. Conversely, patient cells that were enriched for both IFNγ signaling and an invasive phenotype (Mel97 – blue cluster; Mel89 – purple cluster) had the highest PD-L1 levels (**Fig 5F**). This observation also recapitulates our model predictions that neural crest-like or invasive states are highly likely to have upregulated PD-L1 levels (**Fig 5B, right**). Next, we focused on samples which had only one of the two axes being active: proliferative-invasive transition and IFNγ signaling. Samples with high IFNγ signaling but low invasive scores (Mel103 – brown cluster; Mel 71 – orange cluster) had higher levels of PD-L1 than the ones with high invasive score but with low IFNγ signaling (Mel81 – green cluster) (**Fig 5F**). However, both these scenarios had lower PD-L1 levels compared to the “double positive” high invasive and high IFNγ activity subset (**Fig 5F**). Therefore, our dynamical model can reproduce the origins of PD-L1 heterogeneity in melanoma as a function of contributions from proliferative to invasive transition and IFNγ signaling.

### Exploring the Dual Routes of Therapeutic Resistance in Melanoma: Pre-existing and Induced Heterogeneity Pathways and Varied Trajectories

To better decipher the complex interplay between PD-L1 expression, proliferative-invasive transition, and IFNγ signaling in melanoma under the influence of both targeted therapies and immune checkpoint inhibitor therapies, we analyzed transcriptomic profiles of WM989 cells treated with vemurafenib (a potent, highly selective inhibitor of mutated BRAF V600E) after 48 hours of treatment, and 7 days post development of therapy resistance (GSE97681) (65). We observed that melanoma cells post vemurafenib treatment at both these time-points were more invasive and had enhanced IFNγ signaling activity (**Fig 6A**). This combined increase along both IFNγ signaling axis and proliferative-invasive axis is accompanied by an increase in PD-L1 expression levels. A short-term response (48 hours of treatment with vemurafenib) was found to be sufficient to induce a proliferative to invasive transition (**Fig 6A**). This trajectory of drug resistance was also seen in other studies in other datasets for the same cell line (WM989) treated either treated with vemurafenib (GSE161299) (**Fig S9B**) alone or in combination with pinometostat, a clinical-stage DOT1L inhibitor (GSE161298) (**Fig S9C**).

**Figure 6:**
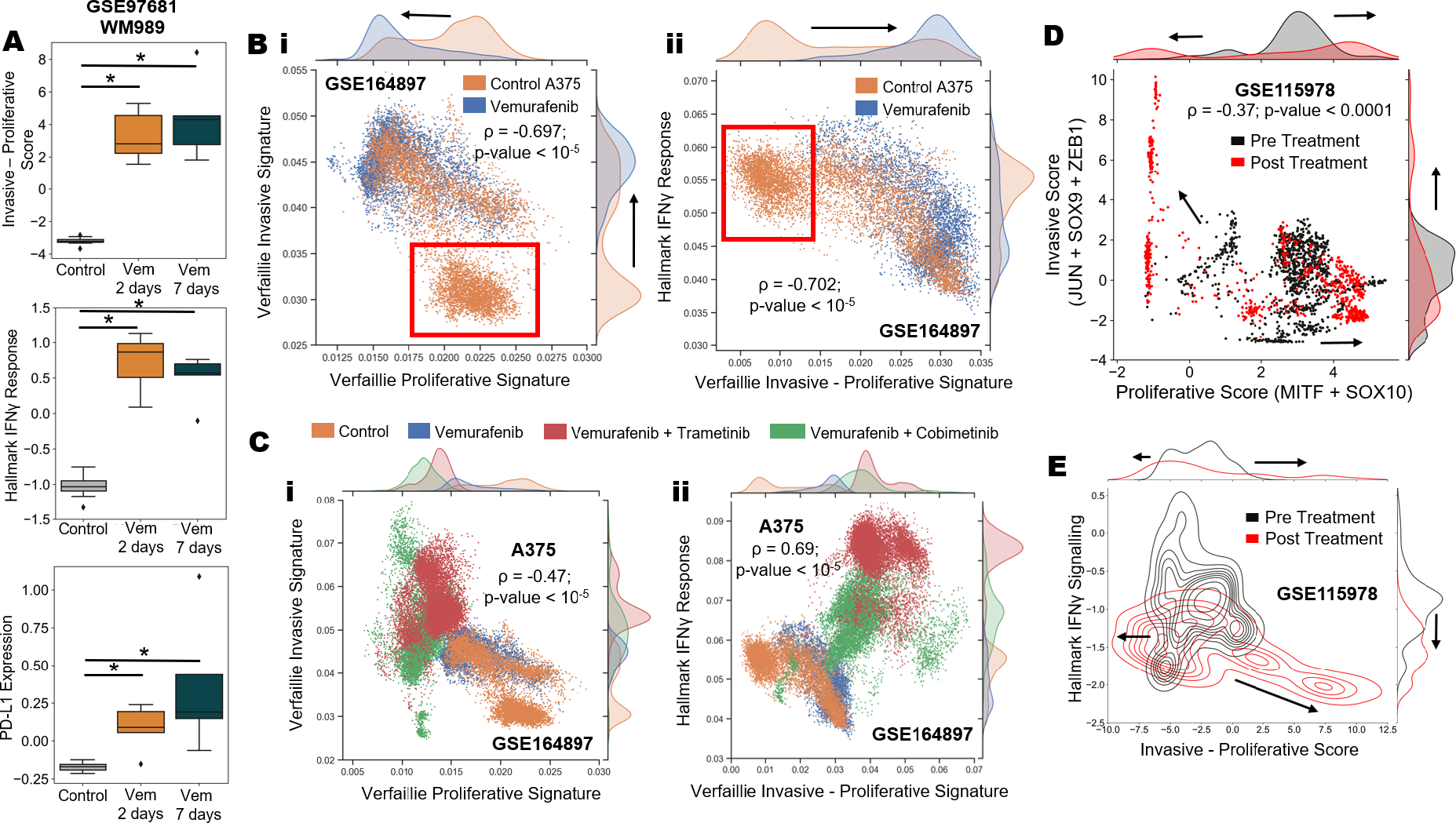
Role of pre-existing and induced heterogeneity in development of drug resistance to both targeted therapy and immuno-therapy. A) Box plot illustrating the changes in proliferative-invasive status (top), IFNγ signaling (middle) and PD-L1 expression levels (bottom), following treatment with vemurafenib for 48 hours and 7 days (top). B) i) Scatterplot of single-cell RNA sequencing (scRNA-seq) data for MEK inhibitor-treated A375 cell line, comparing untreated and vemurafenib-treated conditions, projected on the proliferative-invasive plane based on the imputed expression of Verfaillie signatures. ii) Scatterplot depicting IFNγ signaling levels in the A375 cell line under untreated and vemurafenib-treated conditions. The red box denotes the transcriptional state which was present in untreated but was absent in Vemurafenib resistant clones. C) i) Scatterplots of all A375 cell lines under different MEK inhibitor treatment conditions, projected on proliferative-invasive planes, based on the imputed expression of Verfaillie signatures. ii) Scatterplot illustrating the differences in IFNγ signaling levels under various MEK inhibitor treatment conditions. D) Scatterplot showing immune checkpoint inhibitor pre-treatment (black) and post-treatment (red) single cells projected on the proliferative-invasive plane. Arrows represent hypothesized directions of drift of melanoma cells during adaptation to therapy. E) Contour showing immune checkpoint inhibitor pre-treatment (black) and post treatment (red) single cells projected on the proliferative- invasive spectrum and Hallmark IFNγ signaling. Arrows represent hypothesized directions of drift of melanoma cells due to adaptation to therapy. * denotes a statistically significant difference in levels based on Student’s t-test.

Next, we investigated the single cell transcriptomics profiling of 31,000 BRAF-mutant A375 cells in 4 treatment conditions: untreated/sensitive cells, cells resistant to vemurafenib, cells double-resistant to vemurafenib and cobimetinib, cells double-resistant to vemurafenib and trametinib (GSE164897) (66) We observed that the untreated cells exhibited two distinct subpopulations – one of which exhibited a transcriptional state present only in the untreated cells (**Fig 6B, red box**), and the other overlapped with the state seen in treated cells, indicating pre-existing heterogeneity in terms of varying invasive nature. In the treated population, cells displayed a predominantly more invasive state (**Fig 6B, i**). This change could indicate two possible scenarios that are not necessarily mutually exclusive: a) pre-existing heterogeneity in proliferative-invasive status of cells may have provided more invasive cells with a selective advantage during the evolution of drug resistance to vemurafenib, b) drug treatment induced a subpopulation of cells to alter phenotypes. Furthermore, these cells had lower levels of IFNγ signaling (**Fig 6B, ii**) despite having a higher invasive score. This trend agrees with another instance (GSE108383) of vemurafenib resistant clones of A375 and 451Lu cells characterized by reduced IFNγ signaling compared to their sensitive counterparts (**Fig S9D**). We also observed that a slight increase in PD-L1 levels post-treatment in vemurafenib-resistant cells, and a positive correlation with the invasive status (**Fig S9E**). These results indicated that in some adaptation strategies, the positive association between proliferative to invasive transition and IFNγ signaling can be compromised, while the levels of PD- L1 and the extent of proliferative to invasive transition can increase concurrently.

On the other hand, in case of combinatorial drug treatment, i.e., treatment with BRAF inhibitor vemurafenib in combination with MEK1 inhibitor trametinib or cobimetinib, we observed the emergence of another phenotype that was even more invasive than the vemurafenib-resistant one (**Fig 6C, i**). This state was earlier not observed in untreated samples, and was accompanied by an increase in IFNγ signaling (**Fig 6C, ii**), suggesting an induction to a distinct new transcriptional state not previously present in the pre-treatment setting. Such new attractor states enabled by the induction through combined BRAF/MEK1 inhibition may also contribute to the development of resistance and tumor relapse. Furthermore, the cell-state transition trajectory(ies) during the evolution of resistance to a targeted therapy combination could depend on cell type itself. For example, a combinatorial treatment of MEK and CDK4/6 inhibitors over a prolonged period (∼1 month) promoted four different cell populations (GSE230538) (67), each with distinct cell state trajectories on the proliferative-invasive plane. We observed that the cell lines which were drug-naïve upon treatment over 1, 4 and 33 days may or may not show a reversal to their original proliferative-invasive status during the evolution of resistance. For example, MEL-JUSO and M20 became significantly more proliferative as well as significantly more invasive upon becoming resistant while the cell lines IPC-298 and SK-MEL-30 showed an increase in both proliferative and invasive scores but returned to a proliferative-invasive state like their treatment naïve case at the end of 33 days of drug treatments (**Fig S9F**).

We next investigated changes in transcriptional state along the proliferative-invasive spectrum and IFNγ signaling upon cells becoming resistant to immune checkpoint inhibitors (ICIs). We analyzed single-cell transcriptomics data from patient-derived melanoma cells that are resistant to immune checkpoint inhibitor (GSE115978) therapy (68). Projecting the pre-treatment and post-treatment single-cell profiles along the axes of invasive nature, PD-L1 expression and IFNγ signaling activity, we observed two distinct trajectories that can correspond to adaptive response to immune checkpoint inhibitor therapy. In one trajectory, melanoma cells become more invasive, as shown by a distinct increase in invasive score and a concomitant decrease in proliferative scores (**Fig 6D**). These cells also have reduced IFNγ signaling activity (**Fig 6E**). This trajectory was recently reported in experimental data demonstrating that melanoma patients responding to anti-PD-1 therapy exhibit a more proliferative gene expression signature, while the non-responders were enriched in invasive and neural crest-like phenotypes (69). Another trajectory that cells can follow is cells exhibiting a more proliferative nature, without a noticeable change in IFNγ signaling activity (**Fig 6D, E**). A similar trajectory is reported in response to anti-BRAF therapy, where cells can exhibit a hyperdifferentiated phenotype (70, 71). Finally, we analysed post-treatment samples, and observed diverse trajectories of drug adaptation. We plotted the single-cell RNA-seq data from melanoma patients’ post-therapy on two-dimensional plane of proliferative-invasive score and IFNγ signaling (**Fig S9G, top**). Similar to treatment naïve samples (**Fig 5F**), inter-patient variability dominates over intra-patient variability across both the axes (**Fig S9G**). We observed that the least invasive cells had the highest expression of PD-L1 levels (Mel88, Mel78, Mel98 - orange, blue and pink clusters, respectively). However, these cell clusters were also higher in IFNγ signaling compared to the more invasive clusters, such as Mel110 (purple) and Mel106 (brown), which had relatively lower levels of PD-L1 (**Fig S9G, bottom**). These results indicate that IFNγ signaling levels may be a dominant determinant of PD-L1 levels as compared to the proliferative-invasive status of cells in immune checkpoint inhibitor-resistant melanoma cells. Such co-occurrence of adaptive drug-tolerant states is reflective of how isogenic cells exposed to identical therapeutic stress can diverge towards different “attractors” in the phenotypic landscape, driving non-genetic heterogeneity. Together, our results highlight the importance of both pre-existing heterogeneity and cell-state transition driven heterogeneity along the different trajectories on the IP and IFNγ signaling axes, as two distinct but not necessarily exclusive routes to emergence of drug resistance in cancer cells. However, the exact underlying mechanisms of resistance, and the interplay between these two routes, require further investigation and more mechanistic modeling, indicating the future scope of our study.

Overall, our dynamical model, coupled with single-cell transcriptomic analysis of melanoma samples, indicate that both IFNγ signaling activity and proliferative-invasive transition can govern PD-L1 levels in a combinatorial manner in treatment-naïve conditions and specific cases of resistance to targeted and/or immune checkpoint inhibitor therapies. Moreover, diverse cellular trajectories of adaptive response to anti-BRAF or immune checkpoint inhibitor therapy can alter how proliferative to invasive transition and IFNγ-induced dedifferentiation shape the PD-L1 heterogeneity patterns in a tumor.

## DISCUSSION

Melanoma is highly heterogeneous, with diverse underlying genomic (72, 73) and non-genetic features (74, 75). Due to such extensive heterogeneity, standard targeted inhibitor therapies (BRAF inhibitors: dabrafenib, vemurafenib, encorafenib; MEK inhibitors: trametinib, cobimetinib, binimetinib) are effective only in a subset of patients. While combinatorial therapeutic strategies can prolong median overall survival and median progression-free survival, the acquisition of primary and acquired resistance remains a clinical challenge (76). Similarly, immune checkpoint inhibitors have revolutionized treatment for advanced melanoma, but these treatments are not universally effective due the heterogenous expression of the targeted molecules.

Here, we elucidate how this heterogeneity emerges at a non-genetic level. Our minimalistic model of the interactions among key master regulators of proliferative-invasive transition demonstrates can capture the major phenotypic states common to melanoma cells, and their associated immune-suppressive status (**Fig 7A**). Coupling this core network to PD-L1 and IFNγ signaling provides a mechanistic basis for observed PD-L1 heterogeneity in patients (22). We demonstrate that PD-L1 expression can be regulated by combinatorial influence of IFNγ signaling and proliferative-invasive status of a cell, both through dynamical modeling and through our extensive analysis of bulk and single-cell RNA-sequencing data *in vitro*, *in vivo* and from pre- and post-treatment patient data. Of significance, we show experimentally that the combinatorial induction of proliferative to invasive transition and increase in IFNγ signaling can lead to potentially synergistic increase in PD-L1 levels which is significantly higher than individual induction scenarios (**Fig 7B**). We also reveal distinct cell-fate trajectories rooted in both pre-existing resistant cell states as well as induced resistance scenarios that melanoma cells take to adapt to currently used targeted therapies and immune checkpoint inhibitors (**Fig 7C**).

**Figure 7:**
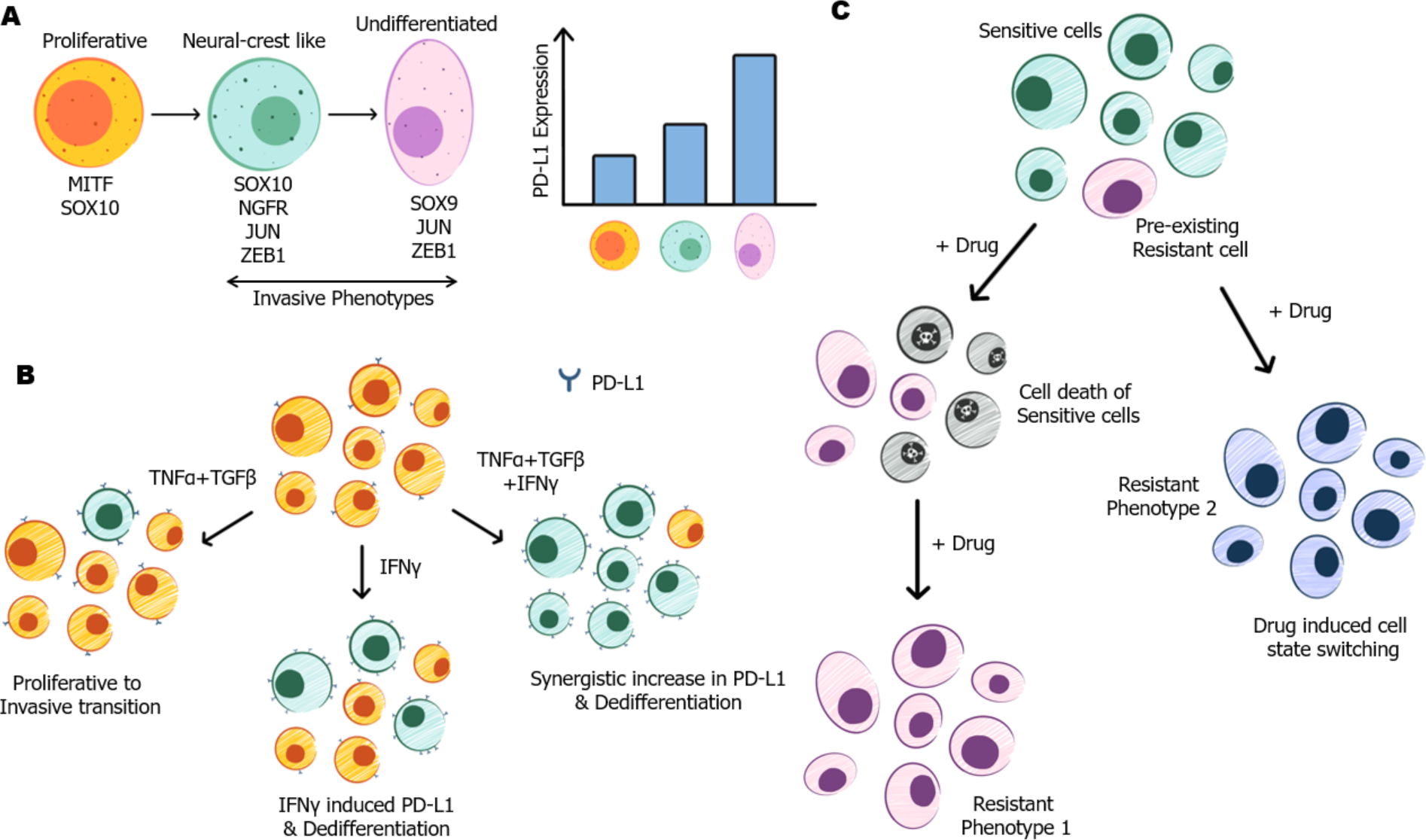
Schematic showing **A)** phenotypic changes during proliferative to invasive transition and associated increase in PD-L1 levels, **B)** Individual and combinatorial effects of proliferative to invasive transition inducers and IFNγ on melanoma cell population and its effects on dedifferentiation and PD-L1 expression, **C)** Distinct paths to adaptation during emergence of resistance to targeted and/or immunotherapies.

Our model, in its current form, has several limitations. First, we did not find any direct evidence in the literature for the existence of an intermediate proliferative/invasive phenotype based on the predicted gene signature. This prediction can be explored by further experiments to identify a MITF^high^/SOX9^high^ state in preclinical and clinical samples. Second, our model could not explain the existence of the hyper-differentiated phenotype, which has been reported in the literature (10). Our model is also currently not capable of explaining how the negative association of proliferative-invasive status and IFNγ signaling manifests in treatment resistant cases and how PD-L1 levels are regulated in such scenarios. Future iterations on network topology are needed to identify the molecular interactions enabling this scenarios. Third, our model is a simplified minimalistic one addressing cell-autonomous behavior, while melanoma cell behavior is affected by tumor microenvironment and immune cells too (77). Future multi-scale models, similar to ones investigating tumor-immune crosstalk (78), can be helpful in delineating how these external signals affect cellular heterogeneity and subsequently the therapeutic response.

Despite these limitations, our *in silico* dynamical model, well-calibrated with *in vitro*, *in vivo,* and patient transcriptomic data both at single-cell and bulk levels, can serve as a platform to both better understand the emergence of phenotypic heterogeneity in melanoma and identify the trajectories of adaptive resistance to existing therapies. The experimental validation of synergistic increase in PD-L1 levels as predicted by the model opens up new avenues for targeting IFNγ signaling as well as proliferative to invasive transition inducing pathways to simultaneously inhibit melanoma metastasis and immune evasion via T cell exhaustion. We envision that our model can be integrated with the patient-calibrated mathematical models (79) to suggest more effective interventions for metastatic melanoma, and to advance the boundaries of precision medicine-based treatment paradigms.

## Author contributions

SS (Seemadri) and SS (Sarthak) performed research, analyzed data and prepared a first draft of manuscript. SD performed in vitro experiments. MKJ designed and supervised research and obtained research funding. JAS analyzed data. MKJ and JAS edited the manuscript.

## MATERIALS AND METHODS

### Gene regulatory network simulations using RACIPE

The dynamics of gene regulatory networks (GRNs) specific to biological processes can be explored extensively by using random circuit perturbation (RACIPE), a computational framework that takes the topology of a core regulatory circuit as an input and generates an ensemble of unbiased circuit models with distinct randomized kinetic parameters and initial conditions (80). The input topology of a regulatory circuit delineates the number of genes (nodes) in the network and regulatory links (activating/inhibitory) connecting them. The input t-node regulatory network is then mathematically modeled as a set of chemical rate-based non-linear ordinary differential equations (ODEs).

For a node T in the network having P_i_ activating and N_j_ inhibiting nodes with incoming edges, the ODE generated by RACIPE to represent the time evolution of concentration of node T is as follows:

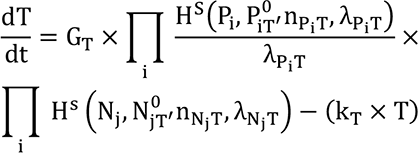

where the terms T, P_i_, and N_j_ are concentrations of nodes at time t (gene expression levels), n is Hill coefficient showing the influence of P_i_ or N_j_ on T, λ is fold change in expression of T, caused by regulatory node P_i_, or N_j_, P^0^_i_ or N^0^_j_ are threshold values of Hill function, G_#_ and k_#_ is production rate and degradation rate of node T respectively. H^S^ (shifted hill function) takes in the activatory/ inhibitory links in account to determine the production rate for the node and is defined by:

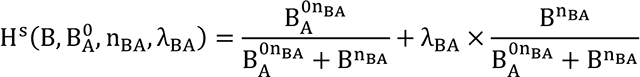

For a particular input topology file, RACIPE generates multiple randomized parameters sets and simulates them over multiple initial conditions to identify steady-state levels of the nodes. Different initial conditions are randomly chosen from a log-uniform distribution ranging from the minimum possible level to the maximum possible level. The parameters are randomized by sampling from their respective biologically relevant pre-defined ranges (Table 1) given below:

**Table 1:** Ranges of randomized parameters in RACIPE.

| Parameter | Minimum Value | Maximum Value |
| --- | --- | --- |
| Production Rate (G) | 1 | 100 |
| Degradation Rate (k) | 0.1 | 1 |
| Inhibition Fold Change ($\lambda^-$) | 0.01 | 1 |
| Activating Fold Change ($\lambda^+$) | 1 | 100 |
| Hill's Coefficient (n) | 1 | 6 |
| Threshold | Half-Functional Rule: ensures that each regulatory link in the network has approximately 50% chance to be functional |  |

The “half-functional” rule is employed to numerically estimate the threshold values in the shifted Hill functions, and it ensures that each regulatory link in the network has approximately 50% chance to be functional, in the ensemble of parameter sets simulated. For example, in the case that gene A regulates gene B, all the threshold parameters are selected in such a way that, for the RACIPE ensemble, the level of A at the steady states has roughly 50% chance to be above and 50% chance to be below its threshold level. If the threshold level is too large or too small, the regulatory link is either not functional most of the time or constitutively active, thereby changing the effective circuit topology, and limiting a comprehensive understanding of the circuit function. Thus, the “half-functional rule” estimates the range of the threshold levels by a mean-field approximation and using it to randomly sample the threshold parameters. For a regulatory link from gene A (regulator) to gene B (target), the threshold level AB^0^ can be estimated as follows. First, the range of expression of gene A is estimated without considering any of its regulators. The A level without regulation satisfies,

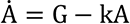

By randomizing both G and k by pre-specified limits (Table 1), an ensemble of random models is generated, from which the distribution of the steady state levels of gene A is obtained. To meet the half-functional rule, the median of the threshold level should be chosen to be the median of this distribution. When gene A is regulated by other genes (i.e., its upstream regulators), its median threshold level is estimated by taking A’s regulators into account, and with assumption that the levels of all these regulators (e.g., gene B, C etc.) follow the same distribution as an isolated gene.

The possible steady state solutions are obtained by solving the coupled ODEs using Euler method. RACIPE provides steady state gene expression data and their characteristic kinetic parameters as an output. The obtained steady state solutions are then Z-normalized.

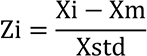

where Xm and Xstd are the mean and standard deviation for a given node, calculated by considering the gene expression values of that node across all its steady states.

Simulations for all the networks were performed in triplicates, with 10,000 parameter sets per replicate and 100 initial conditions for each parameter set. The robust dynamical features of the input network can then be statistically analyzed from the output data using several tools like hierarchical clustering and principal component analysis (PCA).

Networks for the proliferative to invasive transition and its coupling with PD-L1 and IFNγ signaling were mathematically modeled using the above procedure. The perturbation analysis of modulating the gene expression level of a given node, were also carried out using RACIPE by either overexpressing (OE) or down-expressing (DE) the said node by 10-fold. All codes for the RACIPE analysis are uploaded to GitHub and are accessible with the link https://github.com/seemadri/PD-L1_Heterogeneity_Melanoma

### Transcriptomic data analysis – Bulk RNA Seq and Microarray datasets

Publicly available processed microarray and bulk RNA-seq datasets were obtained for melanoma specific datasets (**Table S1**) from GEO as well as from CCLE (skin cell lines) and TCGA-SKCM (patient data). To estimate the activity scores for a pathway/gene list, we used ssGSEA functionality of the python module gseapy (81).

### Gene lists and pathways for activity estimation

To classify the ensemble of steady states into cellular phenotypes more systematically, we defined two scores: proliferative score and invasive score. The proliferative score is calculated by taking the summation of z-normalized expression values of MITF and SOX10, the known master regulators of proliferative phenotype in melanoma. The invasive score is calculated by summing the z-normalized expression values of genes that are known to be associated with cell invasion and metastasis: SOX9, JUN and ZEB1.

Proliferative score = Σ (z-normalized expression of MITF + z-normalized expression of SOX10)

Invasive score = Σ (z-normalized expression of SOX9 + z-normalized expression of JUN + z-normalized expression of ZEB1)

Invasive - Proliferative score = Invasive score – Proliferative score (as defined above)

In the case of including TCF4 and IRF1 in the invasive group of genes, the invasive score and the Invasive - Proliferative score are modified as follows:

Invasive score = Σ (z-normalized expression of SOX9 + z-normalized expression of JUN + z-normalized expression of ZEB1 + z-normalized expression of TCF4 + z-normalized expression of IRF1)

Invasive - Proliferative score = Invasive score – Proliferative score (as defined above)

The signatures for invasive and proliferative nature of melanoma were obtained from Verfaillie et al. (2015) (43). The neural-crest like signature for estimating the neural crest like nature of samples/cells were obtained from Tsoi et al. (2018) (41) and Rambow et al. (2018) (10). The regulon lists for the transcription factors MITF, JUN, SOX10, SOX9 and ZEB1 were obtained from the study by Garcia-Alonso et al. (2019) (82). The hallmark pathways for IFNγ signaling and TNFα signaling were obtained from the list of hallmark pathways from MSigDB (83).

### Transcriptomics data analysis – Single cell RNA-seq datasets

Publicly available count matrices for single cell melanoma datasets for cell lines as well as patient derived lines/samples were downloaded from GEO. Imputation of the gene expression matrices were performed using MAGIC algorithm (84). Activity scores for pathways/gene lists were computed on the imputed gene expression values using AUCell (85).

### In-vitro experiments

Melanoma cells (A375, MALME 3M, SKMEL-5) were seeded at a concentration of 4x10^4^ per well in a 6 well plate format. Cells were then pre-treated with either TNFa (100ng/mL)/TGFb (20ng/mL) or IFN-gamma (10 ng/mL) for 48 hr. mRNA was extracted according to manufacturer’s protocol (Zymo Research) and 500 ng of mRNA was reverse transcribed into cDNA (High-Capacity cDNA Reverse Transcription Kits). Finally, mRNA levels of c-Jun, Zeb1, Ngfr2, MITF and PDL1 were measured and normalized with respect to GAPDH.

### Statistical analysis

Spearman correlation coefficients and corresponding significance levels (p-values) were computed to estimate the degree of association between gene expression/scores and/or activity scores of pathways. Paired/Unpaired students t-tests as well as the corresponding significance values were performed to assess the changes in mean levels of gene expressions and/or pathway activities.

## SUPPLEMENTARY FIGURES

**Figure S1:**
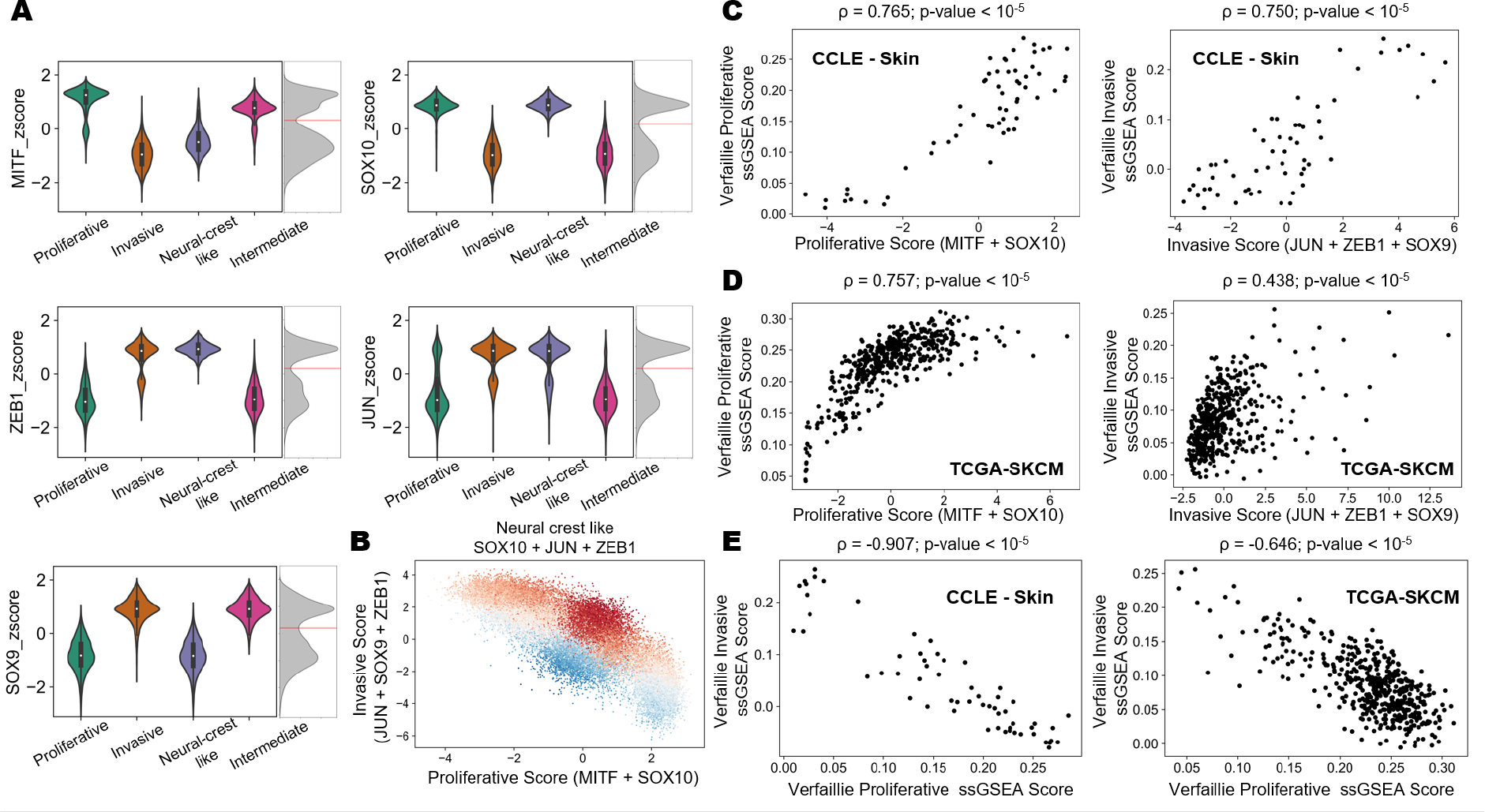
Characteristic gene expression profiles of master regulators for different phenotypes and benchmarking against Verfaillie scores. A) Violin plots of z-normalized steady state gene expression values of MITF, SOX10, ZEB1, JUN and SOX9 grouped by cluster labels obtained from hierarchical clustering. Kernel density estimates for steady state expression of master regulators showing bimodality, partitioned by red line. B) Scatter plot showing the spread of steady state solutions with Proliferative score on x-axis and Invasive score on y-axis. The steady states have been colored by Neural crest like scores. C) Scatter plot comparing five gene based proliferative and invasive score against ssGSEA score for Verfaillie proliferative (left) and invasive gene signatures (right) in CCLE group of skin cancer cell lines. D) Scatter plot comparing five gene based proliferative and invasive score against ssGSEA score for Verfaillie proliferative (left) and invasive gene signatures (right) in TCGA cohort of SKCM patients. E) Scatter plot showing association between Verfaillie proliferative and invasive ssGSEA scores for CCLE group of skin cancer cell lines (left) and TCGA cohort of SKCM patients (right).

**Figure S2:**
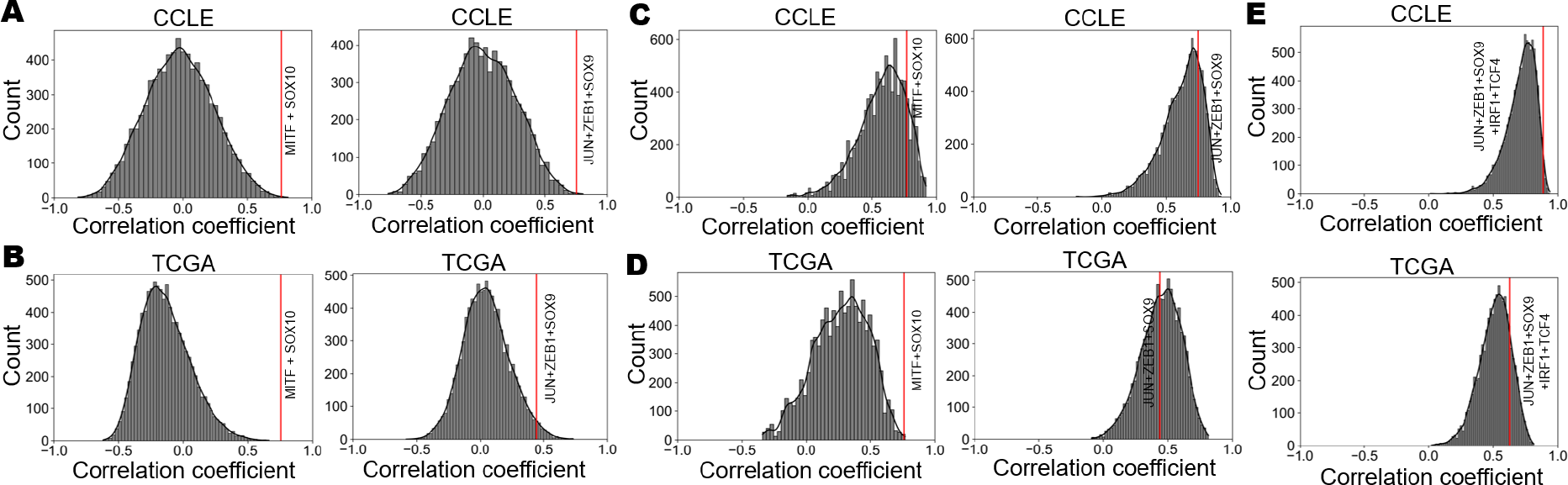
Assessing random gene combination correlations with Verfaillie signatures. **A)** Frequency distribution of correlation coefficients for random combinations of any 2 (left) or 3 (right) transcription factors with Verfaillie proliferative (left) and invasive (right) gene signature ssGSEA scores in the CCLE skin cancer cell line group. **B)** Same as A) but for TCGA SKCM patient cohort. **C)** Frequency distribution of correlation coefficients for random combinations of 2 (left) or 3 (right) transcription factors chosen from within the Verfaillie proliferative (left) and invasive (right) signatures in the CCLE skin cancer cell line group. **D)** Same as C) but for TCGA SKCM patient cohort. Red line indicates the correlation of proliferative score (MITF+SOX10) with the Verfaillie proliferative signature (left) and the correlation of invasive score (SOX9+ZEB1+JUN) with the Verfaillie invasive score in A, B, C, and D. **E)** Frequency distribution of correlation coefficients for random combinations of 5 transcription factors chosen from within the Verifaillie invasive signature in CCLE skin cancer cell line group (top) and TCGA SKCM patient cohort (bottom). The red line represents the correlation of the refined invasive score (SOX9+ZEB1+JUN+IRF1+TCF4) with the Verfaillie invasive score.

**Figure S3:**
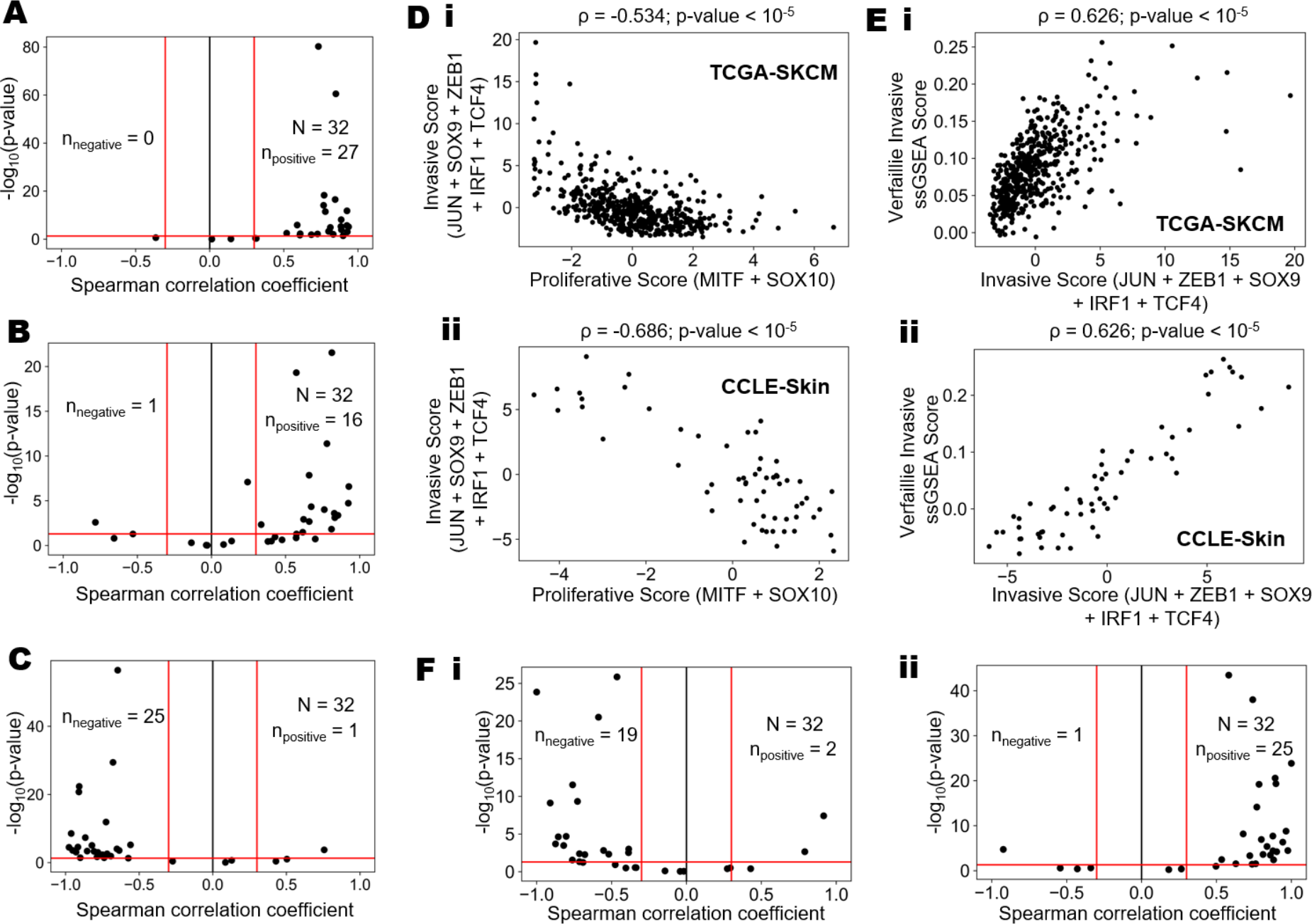
Meta-analysis of melanoma datasets. **A)** Volcano plots showing correlation of two-gene based proliferative score with Verfaillie proliferative gene signature based ssGSEA score **B)** Volcano plots showing correlation of three-gene based invasive score with Verfaillie invasive gene signature based ssGSEA score **C)** Volcano plots showing correlation of Verfaillie proliferative ssGSEA score with Verfaillie invasive ssGSEA score, for all bulk transcriptomics datasets. nnegative and npositive denote the number of datasets (out of 32) that are correlated negatively (Spearman correlation coefficient < -0.3; p-value < 0.05) and positively (Spearman correlation coefficient > 0.3; p-value < 0.05). **D)** Scatterplot showing association between the proliferative (MITF+SOX10) and invasive (SOX9+ZEB1+JUN+IRF1+TCF4) scores for clinical samples from i) TCGA cohort of SKCM patients ii) CCLE-skin cell lines. **E)** Scatter plot comparing five gene based invasive score against ssGSEA based Verfaillie invasive score in i) TCGA cohort of SKCM patients ii) CCLE-skin cell lines. **F)** Volcano plot for the results of meta-analysis of melanoma datasets, accounting for the associations between the i) proliferative (MITF+SOX10) and invasive scores (SOX9+ZEB1+JUN+IRF1+TCF4) ii) invasive scores (SOX9+ZEB1+JUN+IRF1+TCF4) with Verfaillie invasive gene signature based ssGSEA score; nnegative and npositive denote the number of datasets (out of 32) that are correlated negatively (Spearman correlation coefficient < -0.3; p-value < 0.05) and positively (Spearman correlation coefficient > 0.3; p-value < 0.05)

**Figure S4:**
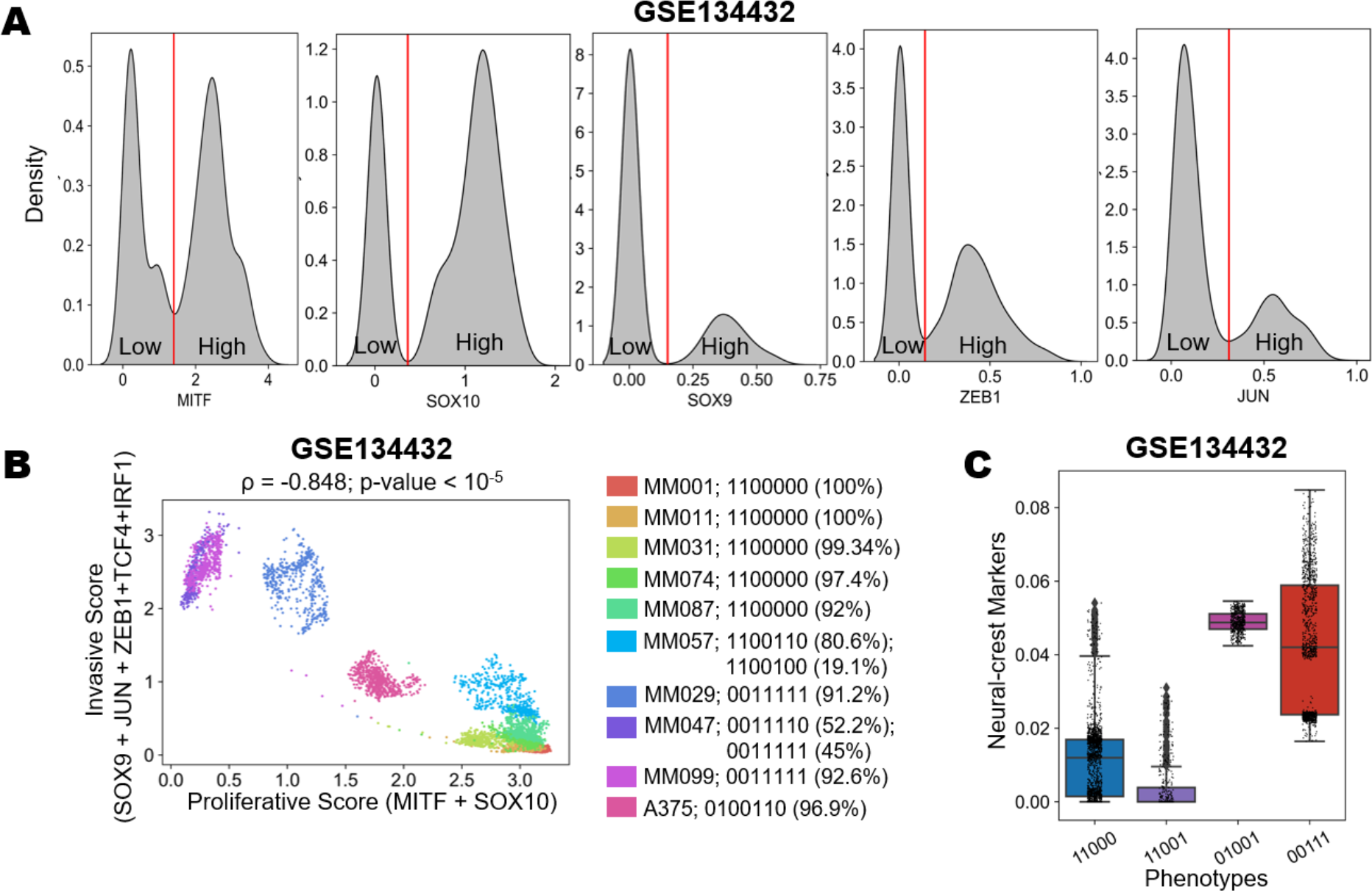
Single-cell transcriptomics data analyses. **A)** Density distributions for expression of master regulators- MITF, SOX10, SOX9, ZEB1, JUN as see in the single cell RNA-seq dataset GSE134432.The red line partitions the expression profiles of these genes to high and low levels at the major minima of each distribution. **B)** Scatterplot of single cell RNAseq data showing each cell of each cell line projected on a proliferative-invasive plane define by proliferative (MITF+SOX10) and invasive (SOX9+ZEB1+JUN+IRF1+TCF4) imputed scores. **C)**Boxplots of cells categorized by the dominant binary phenotypes based on the five gene signature and ssGSEA scores based on Neural-crest markers (GSE134432).

**Figure S5:**
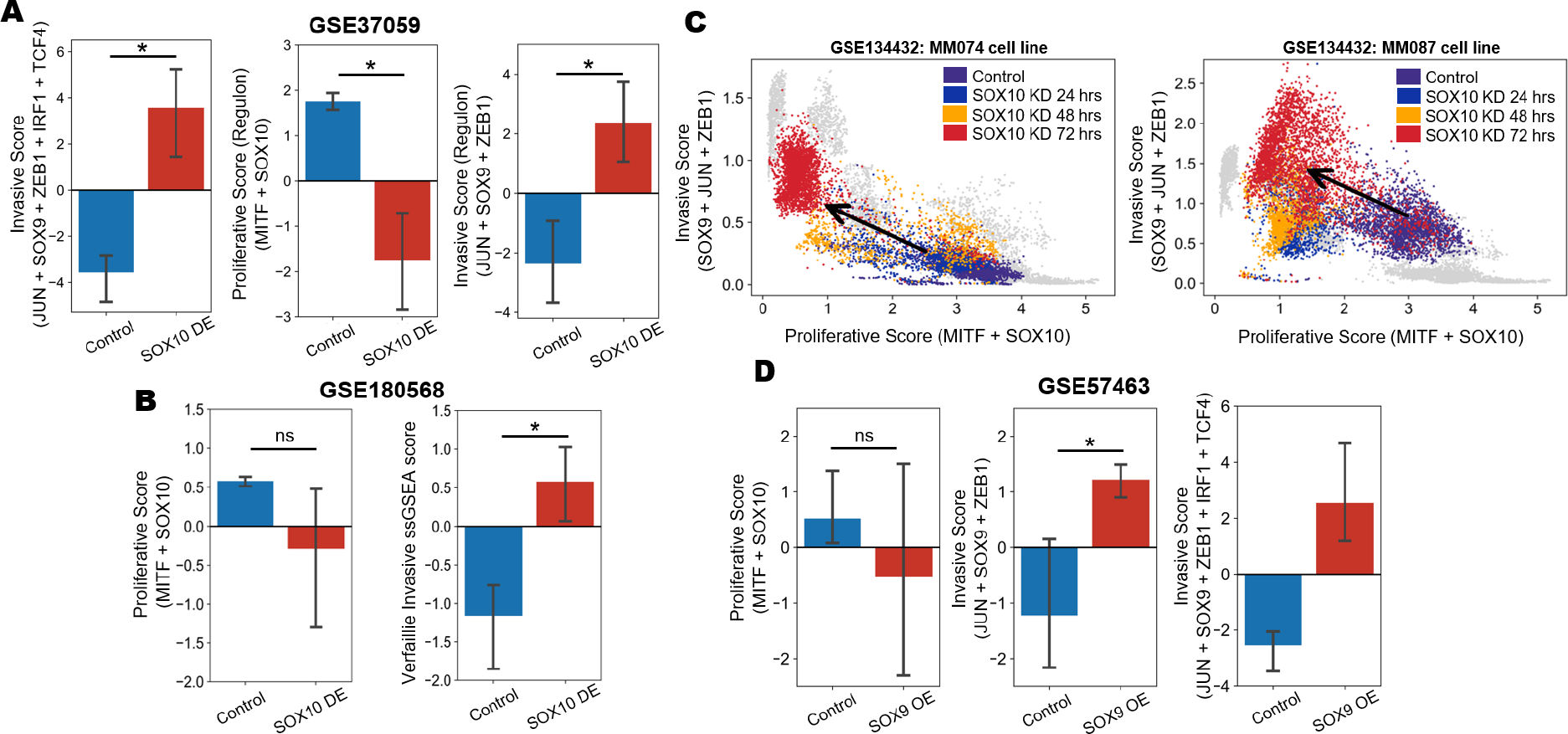
Transcriptomics data in support of transitions along the proliferative-invasive axis upon experimental perturbations. **A)** Bar plots of the experimentally observed significant changes (demarcated by *) in the five gene based invasive scores with TCF4 and IRF1 (left), in the proliferative (defined as the sum of z- normalized ssGSEA scores of MITF and SOX10 regulons) (middle) and invasive scores (defined as the sum of z- normalized ssGSEA scores of JUN, ZEB1 and SOX9 regulons) (right) upon SOX10 down expression in comparison to control case (GSE37059). **B)** Bar plots showing changes in proliferative (left) and invasive (right) scores upon SOX10 down expression (GSE180568). **C)** Scatter plots showing the spread of cells of MM074 (left) and MM087 (right) cell line as they transition along the proliferative-invasive 2D plane over a period of 72 hours of SOX10 siRNA treatment. **D)** Bar plots showing experimentally observed changes in the proliferative score (left) and three- gene based invasive score (middle) and refined five-gene based (right) upon SOX9 over expression (GSE57463).

**Figure S6:**
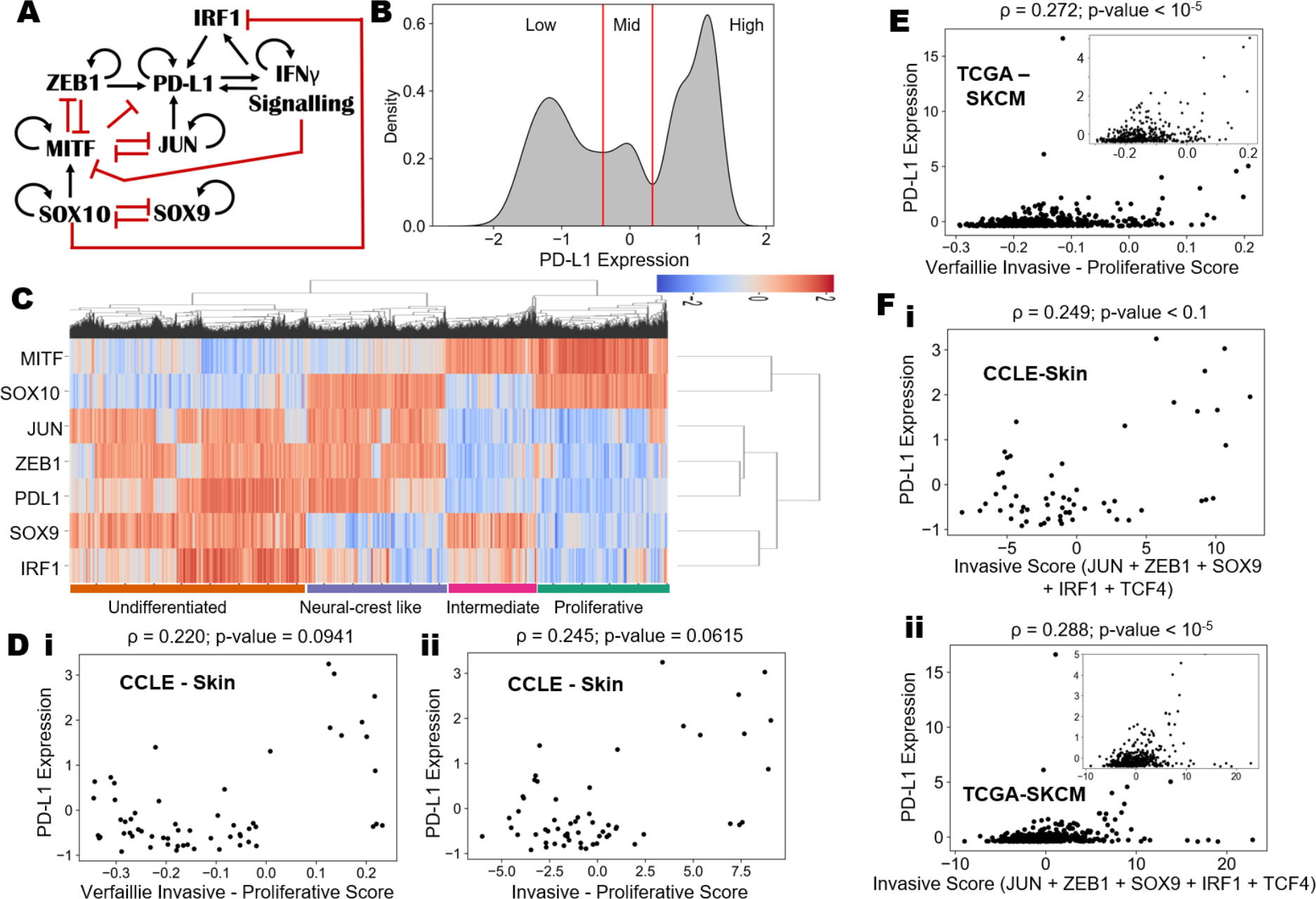
Refining the regulatory network with IRF1 and relation of PD-L1 expression with refined invasive score. **A)** Enhanced gene regulatory network incorporating IRF1 into the previous circuit. **B)** Density histogram of PD-L1 expression fitted with kernel density estimate showing a trimodal distribution. Red lines show the partition between PD-L1 expression levels being high, mid, and low. **C)** Hierarchically clustered heatmap of simulated steady states permitted by the new gene regulatory network and qualitative classification of the four emerging cell states. The simulated four phenotypes have been labelled. **D)** Scatterplot showing associations between the i) Verfaillie Invasive – Proliferative scores and ii) 5 gene signature based invasive – proliferative scores with PD-L1 levels. **E)** Scatterplot showing associations between the Verfaillie Invasive – Proliferative scores with PD-L1 levels in the TCGA cohort of melanoma patients. **F)** Scatterplot showing the association of invasive score with TCF4 and IRF1 and PD-L1 expression for i) CCLE group of skin cancer cell lines ii) TCGA cohort of SKCM patients.

**Figure S7:**
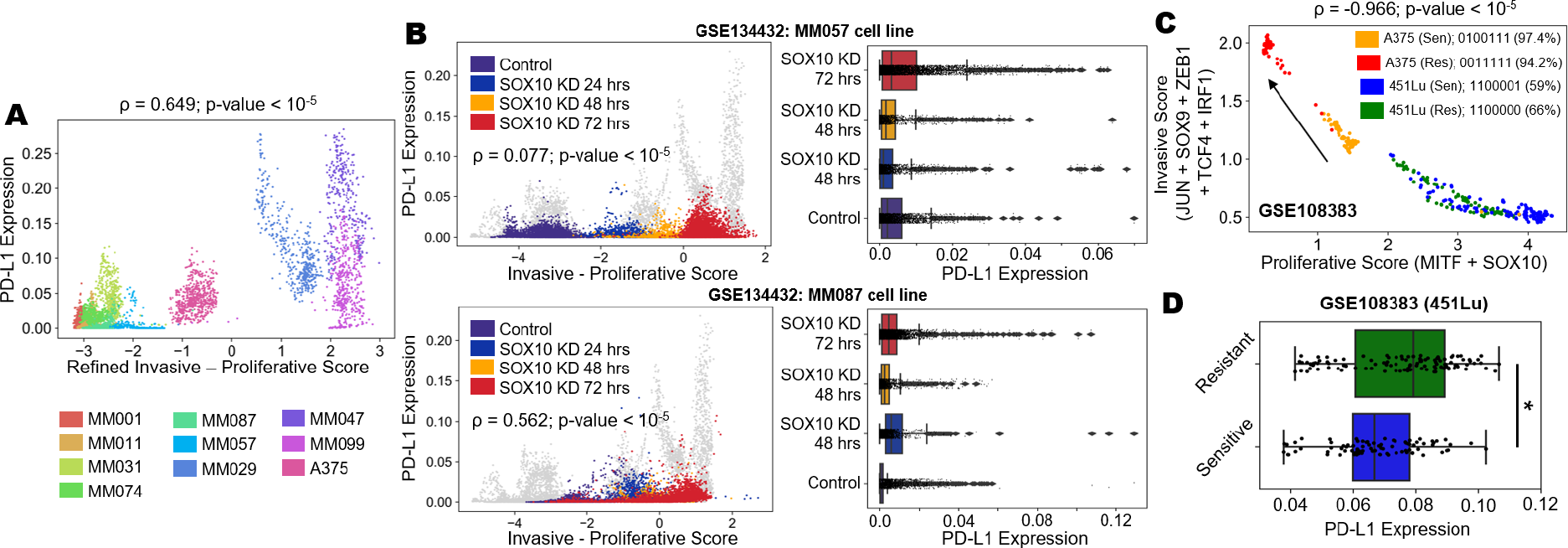
Associations between PD-L1 levels and proliferative-invasive nature of melanoma cells. **A)** Scatterplot showing the association of PD-L1 expression with the new (added IRF1 and TCF4) invasive-proliferative score axis in GSE134432. **B)** Scatter plot and corresponding boxplots showing changes in PD-L1 levels of SOX10 knockdown cells in MM057 (top) and MM087 (bottom) cell lines as they transition from a proliferative phenotype to an invasive phenotype along 24h, 48h and 72h time course single cell RNA-seq data in comparison to control data (GSE134432). **C)** Scatterplot of single cell RNA-seq data projecting cells of two cell lines – A375 (red and orange corresponding to the resistant and sensitive clones, respectively) and 451Lu (green and blue corresponding to the resistant and sensitive clones, respectively) on the refined proliferative-invasive plane. **D)** Box plot showing differences in PD-L1 levels in the sensitive and resistant clones of 451Lu melanoma cells. * represents a statistically significant difference in the levels based on Student’s t-test.

**Figure S8:**
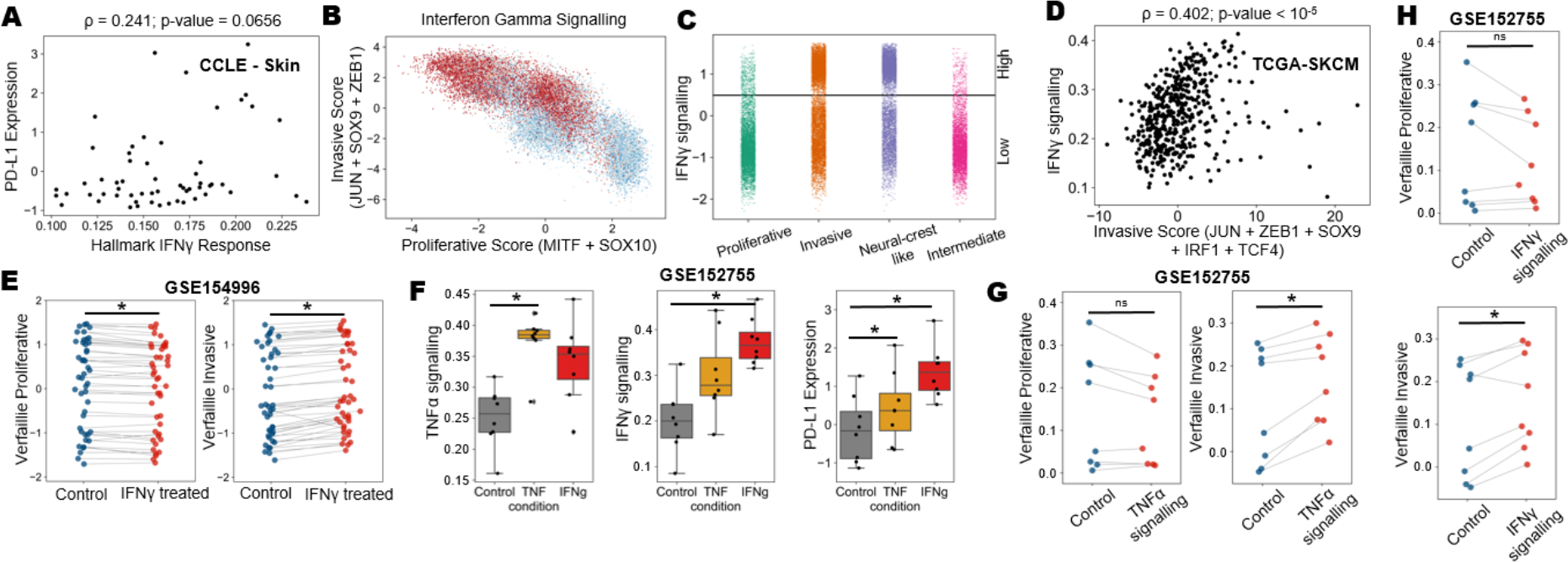
Characterizing association of IFNγ signaling with PD-L1 levels and proliferative invasive nature of melanoma cells. **A)** Scatter plots showing association between Hallmark IFNγ response on x-axis and PD-L1 expression on y-axis for CCLE group of skin cell lines. **B)** Scatter plot showing all the steady states projected onto the proliferative - invasive plane colored based on IFNγ signaling levels. **C)** Strip plot showing the PD-L1 steady state levels for the 4 phenotypes. The horizontal lines mark the stratification of IFNγ signaling levels into low and high regions. **D)** Scatterplot showing the association between IFNγ signaling and invasive score including TCF4 and IRF1 in TCGA cohort of SKCM patients. **E)** Paired plot showing the changes in levels of i) Verfaillie proliferative and ii) Verfaillie Invasive activity levels when wild type melanoma cells are treated with IFNγ. **F)** Boxplot showing levels of Hallmark TNFα, Hallmark IFNγ signalling and PD-L1 levels in 8 melanoma cells treated with either TNF or IFNγ. Paired plot showing the changes in levels of Verfaillie proliferative and Verfaillie Invasive activity levels when wild type melanoma cells are treated with **G)** TNF or **H)** IFNγ. * represents a statistically significant difference in the levels based on a paired Student’s t-test while ns represents a non-significant difference.

**Figure S9:**
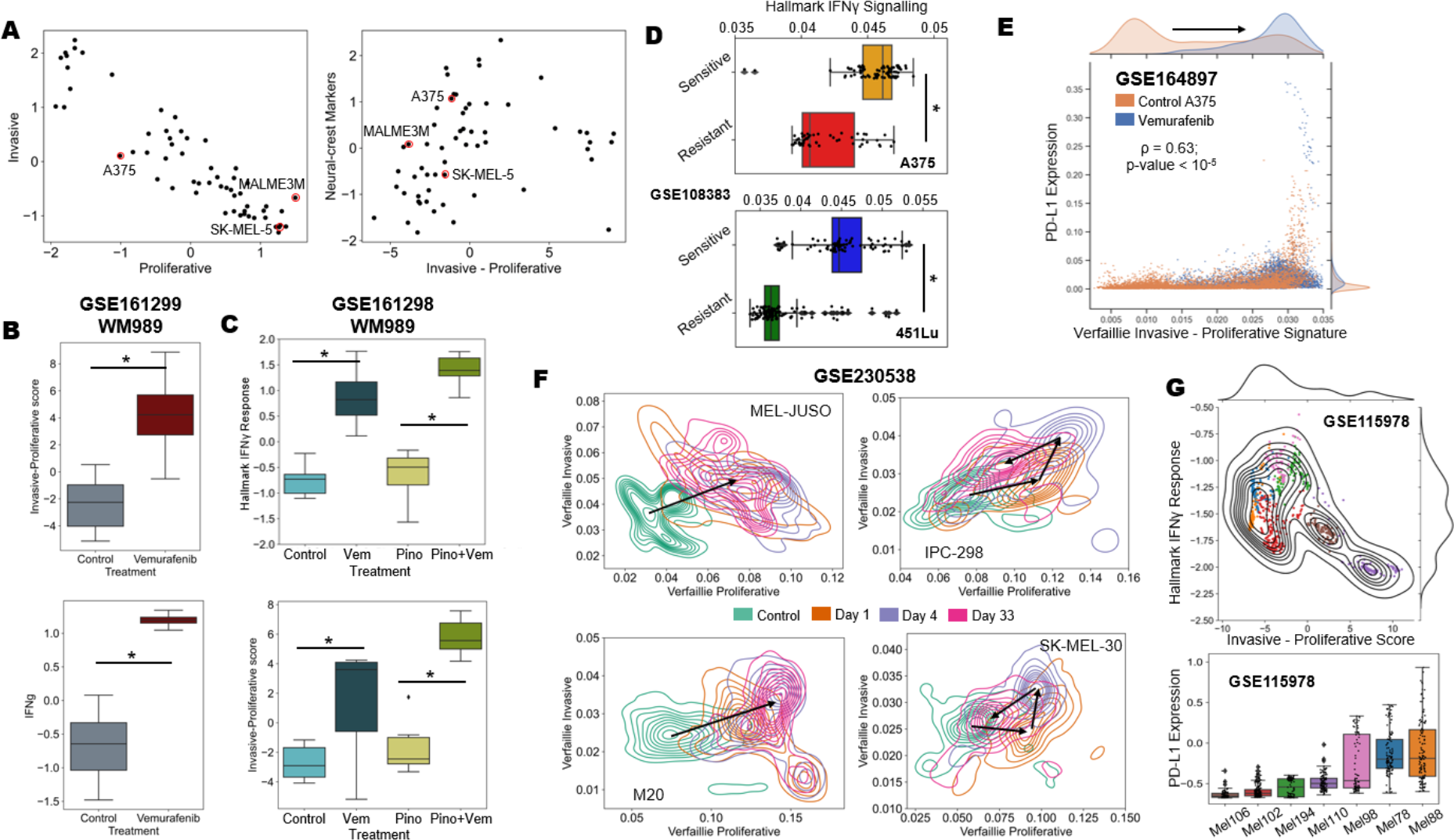
Impact of targeted therapy and immunotherapy on interplay among PD-L1 levels, proliferative to invasive transition and IFNγ signaling. A) Scatterplots showing the association between Verfaillie proliferative and invasive score (left) and proliferative to invasive transition score with Neural crest ssGSEA score (right) in CCLE skin cancer cell line. Cell lines selected for experimentation (SK-MEL-5, MALME-3M, A375) on basis of their proliferative-invasive status are highlighted in red circles. B) Box plots showing changes along the proliferative- invasive axis (top) and IFNγ signaling (bottom) upon vemurafenib treatment (GSE161299). C) Box plots showing changes along the proliferative-invasive axis (top) and IFNγ signaling (bottom) upon vemurafenib treatment alone and in combination with pinometostat (GSE161298). D) Box plots showing changes in Hallmark IFNγ signaling in sensitive and resistant clones of A375 (top) and 451Lu (bottom) melanoma cells. E) Scatterplot showing modest PD-L1 levels upon vemurafenib treatment (GSE164897). F) Contour maps showing transitions on the proliferative- invasive plane after treatment with MEK inhibitors and CDK4/6 inhibitors for 4 different cell lines (GSE230538). G) Scatterplot of single cells from melanoma patients after treatment with immune checkpoint inhibitor therapy projected on the Invasive-Proliferative score and Hallmark IFNγ signaling and their corresponding PD-L1 levels in a patient specific grouping sorted ascending order according to median expression values. Each color in the scatterplot corresponds to a particular patient sample.

